# Tracking coreceptor switch of the transmitted/founder HIV-1 identifies co-evolution of HIV-1 antigenicity, coreceptor usage and CD4 subset targeting

**DOI:** 10.1101/2023.01.21.525033

**Authors:** Manukumar Honnayakanahalli Marichannegowda, Michelle Zemil, Lindsay Wieczorek, Eric Sanders-Buell, Meera Bose, Anne Marie O’Sullivan, David King, Leilani Francisco, Felisa Diaz-Mendez, Saini Setua, Nicolas Chomont, Nittaya Phanuphak, Jintanat Ananworanich, Denise Hsu, Sandhya Vasan, Nelson L. Michael, Leigh Anne Eller, Sodsai Tovanabutra, Yutaka Tagaya, Merlin L. Robb, Victoria R. Polonis, Hongshuo Song

**Affiliations:** Institute of Human Virology, University of Maryland School of Medicine, Baltimore, MD, USA; U.S. Military HIV Research Program, Walter Reed Army Institute of Research, Silver Spring, MD, USA; The Henry M. Jackson Foundation for the Advancement of Military Medicine, Inc, Bethesda, MD, USA; Centre de Recherche du CHUM and Department of Microbiology, Infectiology and Immunology, Université de Montréal, Montréal, Canada; SEARCH, Institute of HIV Research and Innovation, Bangkok, Thailand; Department of Global Health, University of Amsterdam, Amsterdam, Netherlands; Center for Infectious Diseases Research, Walter Reed Army Institute of Research, Silver Spring, MD, USA

## Abstract

The CCR5 (R5) to CXCR4 (X4) coreceptor switch in natural HIV-1 infection is associated with faster progression to AIDS, but the underlying mechanisms remain unclear. The difficulty in capturing the earliest moment of coreceptor switch *in vivo* limits our understanding of this phenomenon. Here, by tracking the evolution of the transmitted/founder (T/F) HIV-1 in a prospective cohort of individuals at risk for HIV-1 infection identified very early in acute infection, we investigated this process with high resolution. The earliest X4 variants evolved from the R5 tropic T/F strains. Strong X4 usage can be conferred by a single mutation. The mutations responsible for coreceptor switch can confer escape to neutralization and drive X4 variants to replicate mainly in the central memory and naïve CD4+ T cells. We propose a novel concept to explain the co-evolution of virus antigenicity and entry tropism termed “escape by shifting”. This concept posits that for viruses with receptor or coreceptor flexibility, entry tropism alteration represents a mechanism of immune evasion *in vivo*.

## Main text

While the majority of the T/F HIV-1 use CCR5 as the coreceptor^1,2^, viruses able to use CXCR4 tend to emerge during the course of infection, a phenomenon termed “coreceptor switch”^3^. Despite the demonstrated correlation between coreceptor switch and faster progression to AIDS, the biological mechanisms underlying this phenomenon have not been clearly elucidated^3–10^. Several fundamental questions remain. First, it is unclear whether the earliest X4 viruses evolved *de novo* from the R5 tropic T/F viruses or are initially transmitted together with the R5 viruses. Second, the precise biological condition driving coreceptor switch is poorly understood. Third, it is uncertain whether the emergence of the X4 variants is a causal factor or a consequence of accelerated CD4 depletion.

Although it is generally considered that the earliest X4 viruses originated *de novo* from the pre-existing R5 viruses^9–12^, the evolutionary pathway leading to coreceptor switch of the T/F HIV-1 has not been documented in natural HIV-1 infection. This limits our understanding of the fundamental aspects of this phenomenon, and consequently, the biological mechanisms remain controversial. It has been hypothesized that the depletion of the memory CD4+ T cells, the major target cells of the R5 HIV-1, could drive the emergence of the X4 viruses in late infection stages^10,13,14^. However, studies on sequential viral isolates showed that the first isolation of the syncytium-inducing (SI) variants usually preceded the onset of CD4 depletion^15–19^. Moreover, the target cells-based hypothesis could not explain why the R5 viruses tend to be predominant in people harboring the X4 viruses, even at late infection stages when the CD4 count was low^20–25^. Another hypothesis is that X4 viruses could be better controlled by the immune system, in particular the neutralizing antibodies (NAbs) and tend to emerge later when the immune system waned^10,12,26^. However, as NAbs with diverse specificities were identified, emerging evidence indicated that X4 variants from both HIV-1 and HIV-2 are more resistant to neutralization, especially to V3 specific NAbs^27–31^. Clearly, further efforts are needed to better understand the mechanisms underlying coreceptor switch, which has direct relevance to HIV-1 pathogenesis as well as therapeutic and cure strategies.

The availability of longitudinal samples soon after HIV-1 acquisition in the RV217 cohort provides a highly unique opportunity to determine the biological mechanisms driving coreceptor switch in natural HIV-1 infection^32^. By tracking the genetic and phenotypic evolution of the T/F HIV-1, we not only studied how the earliest X4 variants originated from the T/F strains, but also characterized the co-evolution of HIV-1 antigenicity, coreceptor usage and CD4 subset targeting underlying the process of coreceptor switch. We propose a novel conceptual model to explain the co-evolution of virus antigenicity and entry tropism termed “escape by shifting”. The central hypothesis is that for viruses with receptor or coreceptor flexibility, entry tropism alteration functions as an evolutionary mechanism of immune evasion to maintain virus entry capability.

## Results

### Identification of participants with coreceptor switch

We determined virus entry tropism for 20 Thai participants in the RV217 cohort at approximately two years after HIV-1 transmission (Supplementary Tables 1, 2). Coreceptor usage assay identified the existence of X4 viruses in four participants (Supplementary Table 2). Of these four, participants 40094, 40436 and 40257 were initially infected by R5-tropic T/F viruses without evidence of superinfection (Extended Data Figs. 1, 2). Thus, their X4 variants most likely evolved from the T/F strains. Participant 40100 was superinfected by an X4 variant at day 262 (Extended Data Fig. 1).

### Coreceptor switch of the T/F HIV-1 is triggered by driver mutations

To capture the earliest moment of coreceptor switch, we tracked the longitudinal evolution of the T/F viruses for participants 40094 and 40257. Both participants were infected by a single T/F virus as demonstrated by single genome amplification (SGA) and deep sequencing (Fig. 1 and Extended Data Fig. 3). This provided an ideal opportunity to elucidate the evolutionary pathway leading to coreceptor switch. The viral population diversified from the T/F virus over time (Fig. 1 and Extended Data Fig. 2). In 40094, the earliest potential X4 variants, which carried two amino acid substitutions at the V3 N301 glycan site (N301K and T303I), appeared at day 422 (Fig. 1). In 40257, the earliest suspected X4 variants, which contained three mutations in the V3 loop (S306R, Q313H and Q327R), emerged at day 264 (Extended Data Fig. 2). In these variants, most of the substitutions outside of the V3 region were descended from the earlier viruses (Fig. 1 and Extended Data Fig. 2). These observations demonstrate that the earliest potential X4 variants evolved *de novo* from the T/F strains.

**Fig. 1.**
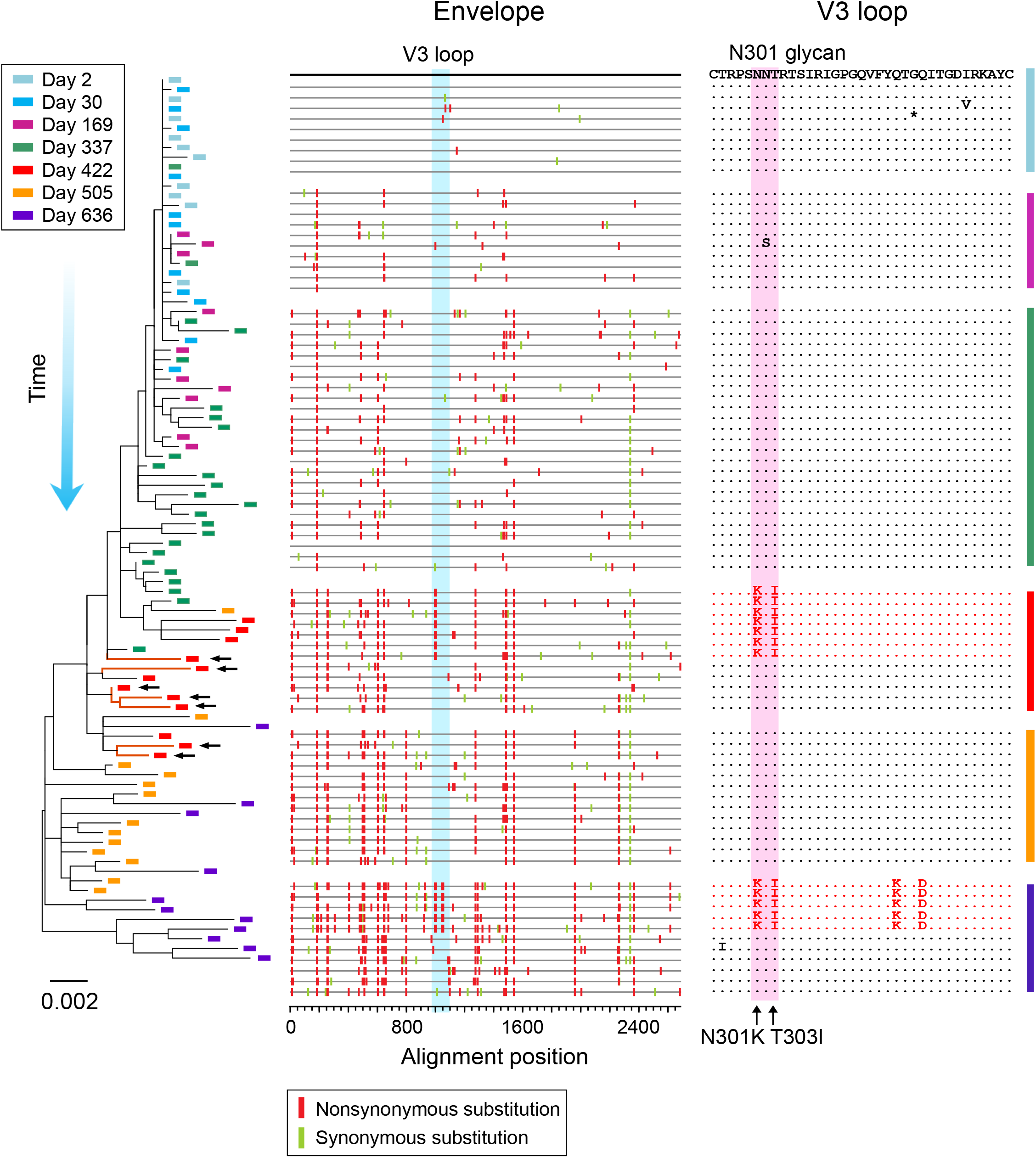
The evolutionary pathway leading to coreceptor switch of the 40094 T/F virus. Longitudinal HIV-1 envelope (*env*) sequences of participant 40094 were obtained by SGA. The evolutionary pathway of the T/F virus is illustrated by using the phylogenetic tree (left) and highlighter plot (middle). Sequences from different time points (days since the first positive test for HIV-1 RNA) are color coded. The phylogenetic tree was constructed using the maximum likelihood method. The earliest X4 viruses are indicated by black arrows. In the highlighter plot, the black line on the top represents the T/F virus, and the red and green tics indicate non-synonymous and synonymous substitutions as compared to the T/F strain. The V3 loop is highlighted in blue. In the V3 amino acids alignment (right), the earliest (founder) X4 sequences are shown in red. The driver mutations responsible for coreceptor switch are indicated by black arrows.

To phenotypically confirm whether these V3 variants were the founder X4 viruses representing the earliest moment of coreceptor switch, we constructed the *env* clones which represented the consensus sequences of these variants. For both participants, the consensus sequences exhibited strong CXCR4 usage and significantly compromised CCR5 tropism in comparison to the cognate T/F virus (15.3-fold and 6.0-fold reduction in 40094 and 40257, respectively) (Fig. 2a-b). We next sought to determine the mutations responsible for coreceptor switch. In 40094, the N301K/T303I mutations, which emerged together at day 422 conferred strong X4 usage on the T/F virus and compromised the efficiency in using R5 (4.3-fold reduction) (Fig. 2a). Analysis of the individual mutations showed that the N301K or T303I mutation were independently sufficient to confer strong X4 usage (Fig. 2a). Reversion of these two mutations back to the T/F sequence nearly abrogated the X4 usage of the founder X4 virus (the “K301N/I303T” mutant) (Fig. 2a). Further reversion of the V1/V2 regions completely abrogated the X4 usage (the “K301N/I303T.V1V2” mutant) (Fig. 2a). In 40257, the three V3 mutations emerged together at day 264 conferred strong X4 usage and significantly compromised CCR5-using ability (8.6-fold reduction) (Fig. 2b). Among them, only the Q313H mutation in the V3 crown was able to confer X4 use (Fig. 2b). Reversion of the V3 mutations back to the T/F sequence completely abrogated the X4 usage of the founder X4 virus (Fig. 2b). These results demonstrate that coreceptor switch of the T/F HIV-1 can be triggered by a single mutation. This finding is in agreement with a previously proposed model based on the structure of CCR5-bound gp120, which hypothesized that HIV-1 coreceptor switch can be accomplished by simple V3 mutations when the V3 loop gains sufficient affinity for CXCR4^33^.

**Fig. 2.**
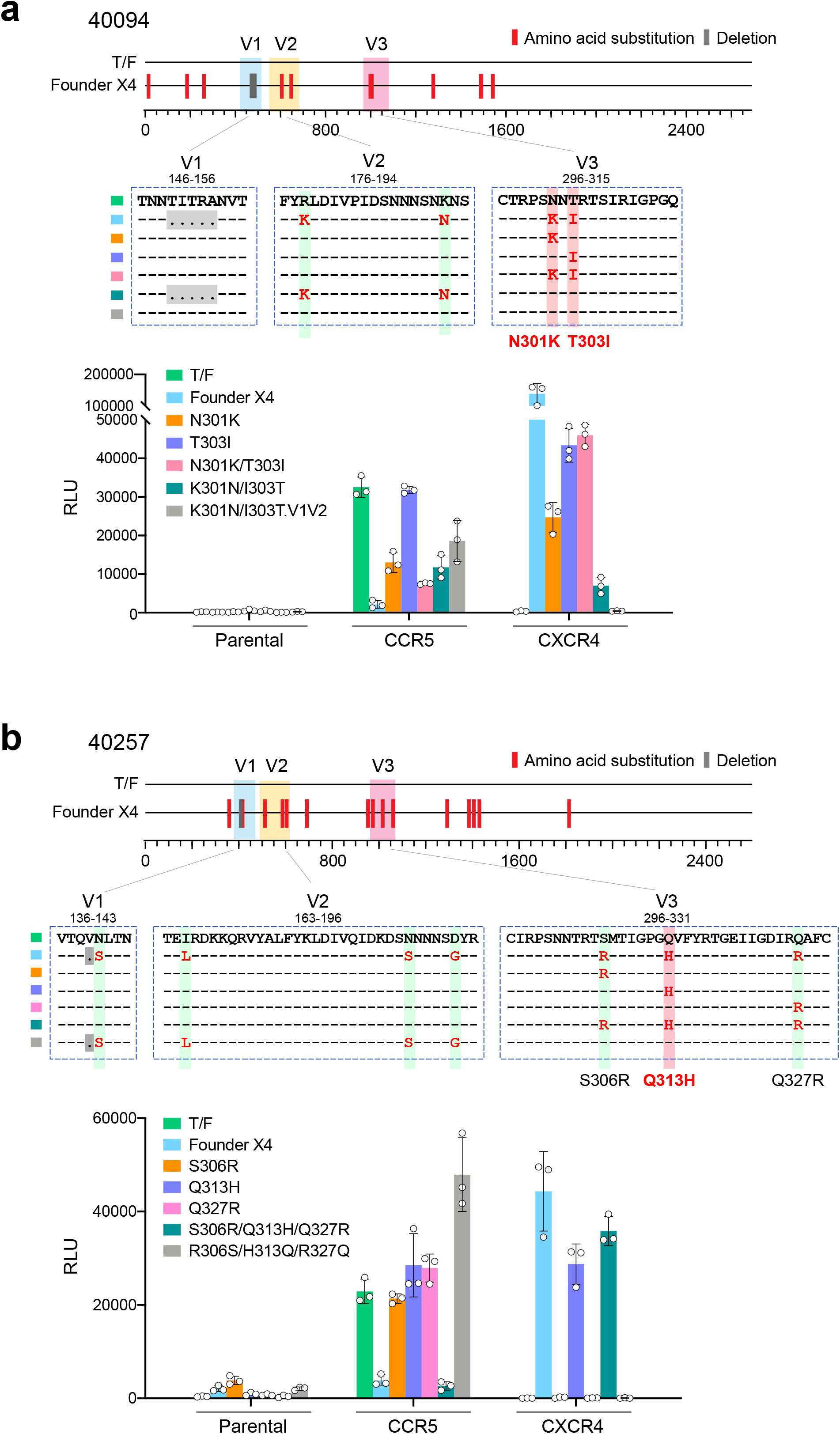
Determination of the driver mutations responsible for coreceptor switch. **a-b**, In participants 40094 (**a**) and 40257 (**b**), the sequence of the founder X4 virus is shown as the consensus sequence of the earliest variants carrying the driver mutations. Amino acid substitutions and deletions in comparison to the T/F strain are indicated by red and gray tics, respectively. The V1, V2 and V3 regions are color coded. Part of the V1, V2 and V3 regions spanning the genetic substitutions are shown. The numbers on the top show the HXB2 locations of the start and the end of each fragment. The driver mutations responsible for coreceptor switch are shaded in red. Different genetic mutants are color coded. Coreceptor usage of the T/F virus, the founder X4 virus, as well as each mutant was determined in NP2 cell lines. All experiments were performed in triplicate and the error bar shows the standard deviation (SD).

To gain a better insight into the evolutionary origin of the founder X4 viruses, we performed deep sequencing on longitudinal plasma samples and tracked the evolution of the variants carrying the driver mutations. In both participants, these mutations could be detected as random mutations at low frequency during acute infection (Extended Data Figs. 4, 5). Similar to the majority of variants detected at acute infection, the variants carrying the X4 usage mutations only differed from the T/F virus by a single base initially (Extended Data Figs. 3–5). Tracking their evolutionary path showed that they were evolving over time, as additional mutations appeared, some of which became detectable by SGA at subsequent time points (Extended Data Figs. 4, 5). However, the X4 driver mutations persisted at low frequency until they were strongly selected for and became detectable by SGA within a short period of time (within 85 and 75 days in 40094 and 40257, respectively). This “deep evolutionary pathway” confirmed that the founder X4 viruses evolved from the T/F virus in a stepwise manner. It also provided direct evidence that coreceptor switch of the T/F virus was likely driven by strong positive selection favoring these V3 mutations.

### Driver mutations confer escape to neutralization

Previous studies showed that X4-using HIV-1 are more resistant to V3 specific NAbs^27,28^. Indeed, both the N301 glycan site and the V3 crown are major targets of the V3 specific NAbs^27,34–36^. These data raised the possibility that the mutations responsible for coreceptor switch could be immune escape mutations. Investigation of the kinetics of autologous neutralization activity showed that in both participants, the V3 mutations were strongly selected for when the autologous neutralization activity was increasing to the peak (Fig. 3a). Determination of the neutralization sensitivity of the founder X4 viruses demonstrated that they were immune escape variants resistant to autologous neutralization (ID_50_ < 20 for 40094; ID_50_ = 88 for 40257) (Fig. 3b). We then determined the impact of the V3 mutations on neutralization susceptibility. The N301K/T303I mutations at the N301 glycan sites, when introduced into the 40094 T/F virus, significantly increased the susceptibility to autologous plasma (*P* = 0.038) (Fig. 3b). However, as previously reported, this effect could be due to the exposure of other antigenic epitopes when the N301 glycan was removed^37,38^. In 40257, the S306R/Q313H/Q327R mutations significantly decreased the susceptibility to autologous plasma (*P* = 0.0071) (Fig. 3b). Among them, only the Q313H mutation could significantly reduce the neutralization susceptibility (*P* = 0.023) (Fig. 3b). Thus, the S306R/Q313H/Q327R mutations selected in 40257 were immune escape mutations, and the escape phenotype was most likely conferred by the Q313H mutation responsible for coreceptor switch.

**Fig. 3.**
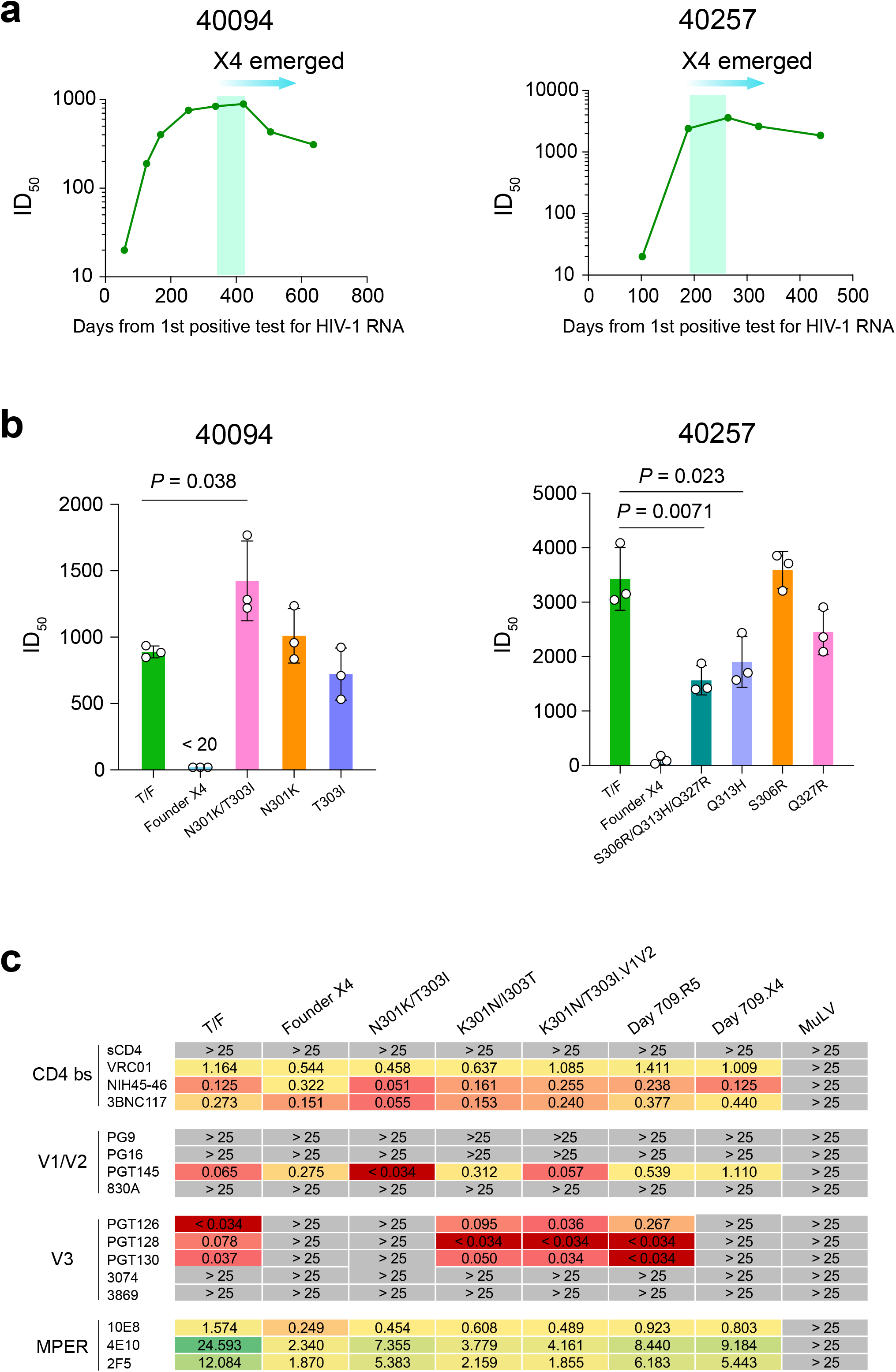
Kinetics of autologous neutralization activity and the impact of driver mutations on neutralization susceptibility. **a**, Kinetics of autologous neutralization activity against the T/F virus. The time frame when earliest X4 viruses emerged is highlighted in green. **b**, Neutralization susceptibility of the founder X4 viruses to autologous plasma and the impact of V3 mutations on neutralization susceptibility. For each participant, plasma samples with the highest neutralization activity against the autologous T/F virus were used for the neutralization assay (day 422 for 40094; day 264 for 40257). The mean titer of three independent neutralization assays is shown and the error bar represents the SD. The statistical significance was determined by using a two-tailed t test. *P* value < 0.05 is considered to be statistically significant. **c**, Neutralization susceptibility of the 40094 T/F virus, the founder X4 virus, as well as different genetic variants to a panel of bNAbs. The IC_50_ (µg/mL) of each bNAb is shown. The murine leukemia virus (MuLV) was used as the negative control.

Genetic substitution at the V3 N301 glycan is a common route to evade the V3 glycan NAbs^34,35,38^. To better understand the impact of the N301K/T303I mutations on neutralization susceptibility, we performed neutralization assays using a panel of broadly neutralizing antibodies (bNAbs) (Fig. 3c). The 40094 T/F viruses can be potently neutralized by the bNAbs targeting the CD4 binding site, the V2 bNAb PGT145, and the V3 glycan bNAbs PGT126, PGT128 and PGT130 (Fig. 3c). However, the founder X4 virus was completely resistant to all V3 glycan bNAbs and had reduced susceptibility to PGT145 (Fig. 3c). Mutations N301K/T303I, when introduced into the 40094 T/F virus, conferred complete escape to all V3 glycan bNAbs (Fig. 3c). In the genetic background of founder X4 virus, reversion of the N301K/T303I mutations to the T/F sequence was sufficient to restore the susceptibility to all V3 glycan bNAbs (Fig. 3c). Further reversion of the V1/V2 regions restored the sensitivity to PGT145 (Fig. 3c). These results demonstrate that the N301K/T303I mutations, which are responsible for coreceptor switch in 40094, are also the exact mutations that confer neutralization escape to V3 glycan bNAbs. Notably, the N301K/T303I mutant had increased sensitivity to PGT145 and the CD4 binding site bNAbs (Fig. 3c), which could be due to the exposure of the corresponding epitopes and might explain its increased sensitivity to autologous plasma (Fig. 3b).

While an X4 virus isolated from day 709 remained resistant to all V3 glycan bNAbs, a co-existing R5 virus remained sensitive to neutralization (Fig. 3c). We further investigated the neutralization sensitivity for another six phenotypically confirmed X4 viruses identified in participants from the RV217 and RV254 cohorts. While most of the T/F viruses in the RV217 Thailand cohort can be neutralized by V3 glycan bNAbs PGT126, PGT128 and PGT130 (Syna Gift et.al., *J Virol*, in press), all X4 viruses identified were fully resistant to bNAbs of this class (Extended Data Fig. 6).

### Driver mutations promote divergence on CD4 subset targeting

The emergence of the driver mutations represents the earliest moment of coreceptor switch. Thus, investigation of the CD4 dynamics before and after this key event provides a unique opportunity to determine the causal relationship between coreceptor switch and accelerated CD4 depletion. Indeed, in both participants, the appearance of the founder X4 virus preceded an accelerated CD4 decline and a continuous increase of the plasma viral load (VL) (Fig. 4). Further analysis of the CD4 subset dynamics in 40094 showed that naïve and central memory (CM) CD4 subsets declined faster than effector memory (EM) and transitional memory (TM) CD4 subsets upon coreceptor switch (Fig. 4a).

**Fig. 4.**
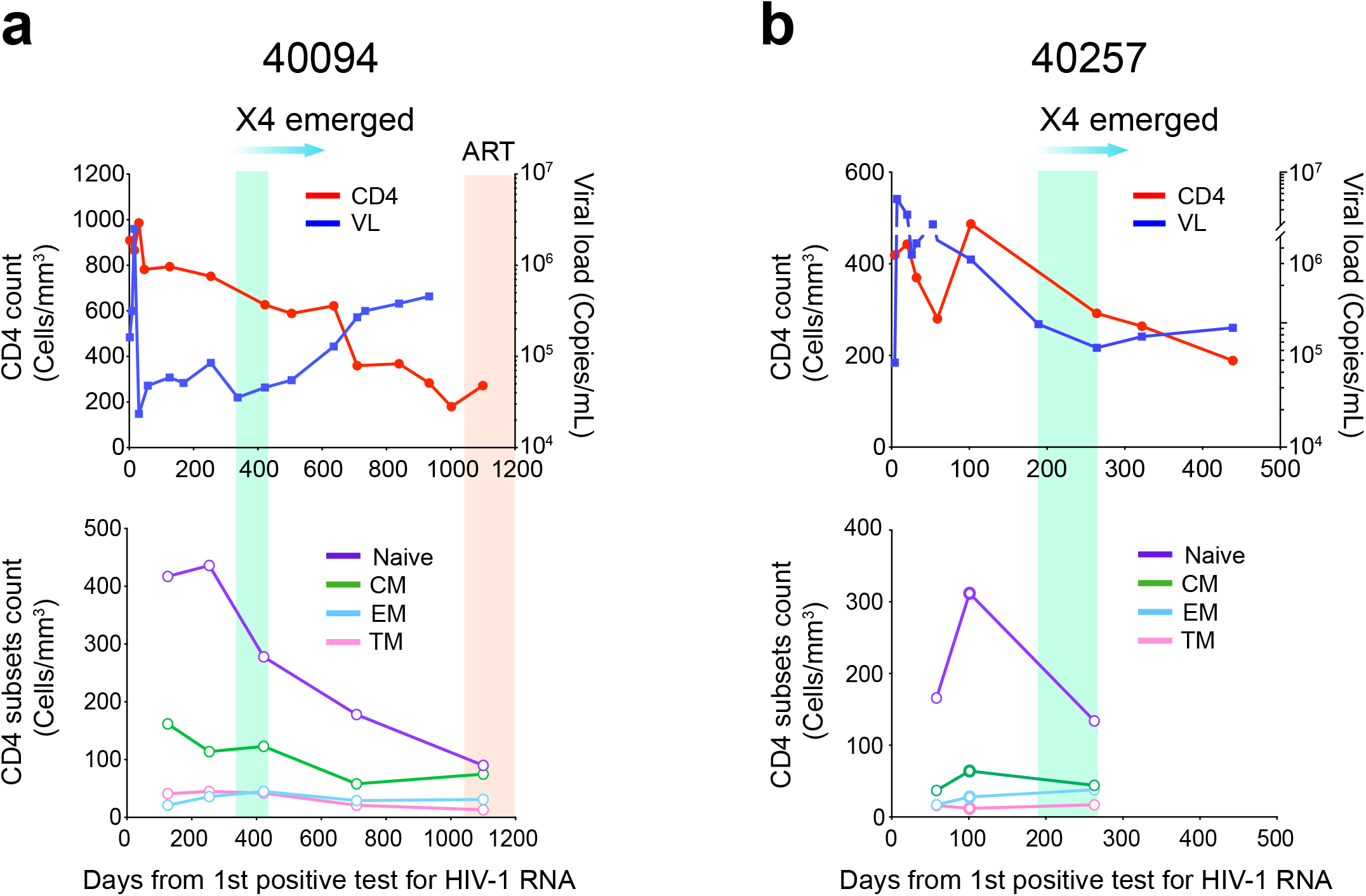
CD4 and viral load dynamics before and after coreceptor switch. **a-b**, Dynamics of CD4 count, plasma viral load (VL) and different CD4 subsets in participants 40094 (**a**) and 40257 (**b**). The time frame when the founder X4 viruses emerged is highlighted in green. In 40094, the time after ART initiation is highlighted in red. The proportion of each CD4 subset was determined by flow cytometry.

To better understand the biological mechanisms, we sequenced the cell associated HIV-1 RNA in each CD4 subset. Sanger sequencing identified a clear compartmentalization of the R5 and the X4 viruses. While the R5 viruses were predominant in the EM and TM CD4 subsets, the emerging X4 viruses were enriched in the CM and naïve CD4 subsets (data not shown). We further sequenced the cell associated HIV-1 RNA by next-generation sequencing and compared their phylogenetic relationship to the viruses in plasma. The analysis revealed distinct cellular origins of the R5 and X4 viruses in plasma. While the R5 viruses mainly originated from the EM, TM and CM subsets, the X4 viruses mainly originated from the naïve and the CM subsets (Fig. 5a). In both participants, the most frequent sequence in the CM subset was X4 tropic, indicating a replication advantage of X4 viruses in the CM cells (Fig. 5a). Consistent with the genetic analysis, the appearance of the founder X4 variants coincided with an emerging cell associated RNA in the naïve cells, which was undetectable earlier, and a substantially increased amount of cell associated RNA (20.5-fold) in the CM cells (Extended Data Fig. 7). We did not measure the dynamic changes of the cell associated RNA in 40257 due to sample availability, but it was also undetectable in the naïve cells before coreceptor switch (Extended Data Fig. 7). Quantification of CCR5 and CXCR4 expression suggested that the compartmentalization was most likely due to the differential expression of the two coreceptors. As previously reported^39,40^, CCR5 expression was relatively high on the EM and the TM subsets, low on the CM subset, and barely detectable on the naïve subset. The CXCR4 expression showed a reciprocal pattern (Fig. 5b). It is important to note that there was no depletion of the TM and EM subsets before coreceptor switch in either participant (Fig. 4). Thus, the origin of the founder X4 virus could not be explained by the depletion of the target cells of the R5 virus.

**Fig. 5.**
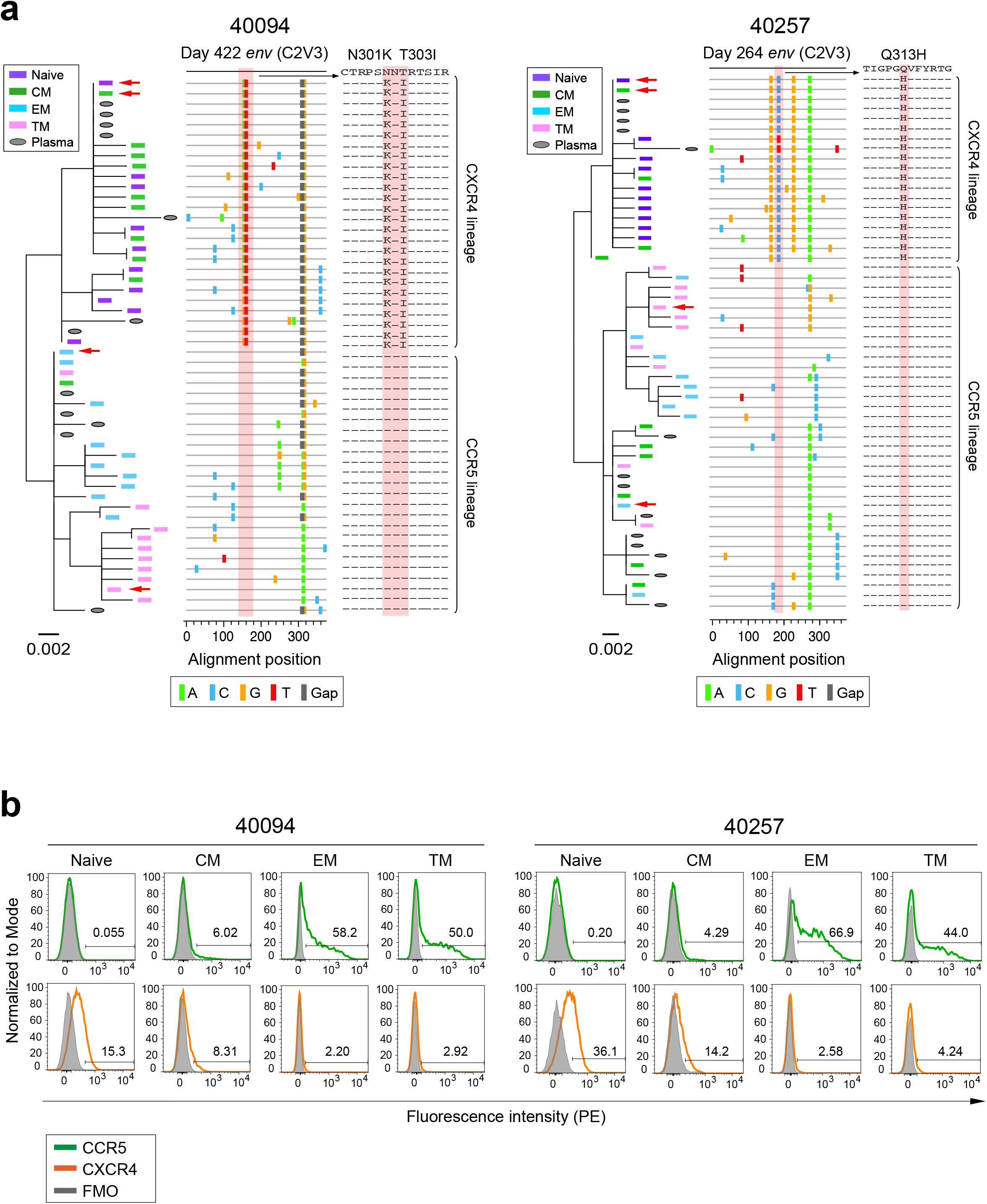
Divergence on CD4 subset targeting of the R5 and X4 HIV-1. **a**, Phylogenetic trees and highlighter plots showing the evolutionary relationship between cell associated HIV-1 RNA and the plasma viruses. The cell associated HIV-1 RNA in each CD4 subset were sequenced by next-generation sequencing using the Illumina MiSeq. Ten unique sequences with the highest frequency in each CD4 subset are shown. Sequences from different CD4 subsets are color coded. The red arrows indicate the most frequent sequence in each subset. In the highlighter plot, the driver mutations responsible for coreceptor switch are shaded in red. **b**, Quantification of CCR5 and CXCR4 expression on each CD4 subset by flow cytometry. Positive cells are shown by percentage in each figure.

To better understand the distinct CD4 subset preferences between R5 and X4 HIV-1, we quantified cell associated RNA for another eleven participants for whom the primary viral isolates showed pure R5 tropism. Cell associated RNA was unexpectedly detected in naïve CD4+ T cells in four participants (Extended Data Fig. 8a). Sequencing of the viruses in naïve cells identified X4 variants in participants 40250 and 40168 (Extended Data Fig. 8b). While we were unable to amplify the *env* region from naïve cells for participants 40577 and 40503, deep sequencing of plasma samples identified low frequency X4 variants in 40577 (Extended Data Fig. 8b). This prompted us to deep sequence the plasma for other participants for whom the PBMC samples were not sufficient to perform cell associated RNA assay. A low frequency of X4 viruses was detected in 40123 (Extended Data Fig. 8b). Participants harboring X4 viruses showed significantly faster CD4 decline (2.7-fold; *P* = 0.0096) (Fig. 6 and Extended Data Fig. 9). Of note, participant 40700, who was infected by an X4 tropic T/F virus (manuscript in preparation), had the fastest CD4 depletion (Extended Data Fig. 9). These results demonstrate distinct CD4 subset preferences of R5 and X4 HIV-1. The data also demonstrate that an X4-using phenotype is a causal factor in driving accelerated CD4 depletion.

**Fig. 6.**
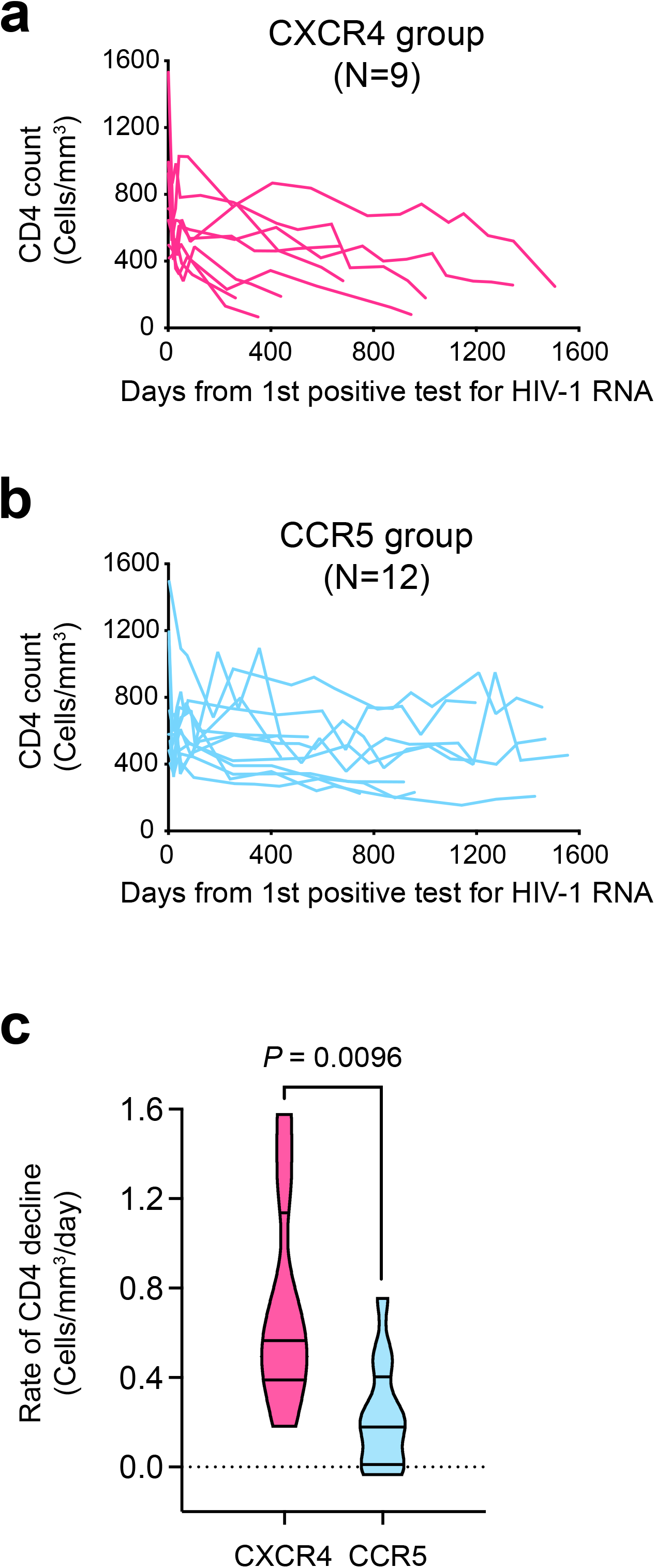
Significantly faster CD4 decline in participants harboring X4 viruses. **a**, CD4 dynamics of participants harboring X4 viruses. **b**, CD4 dynamics of participants without the evidence of harboring X4 viruses. The rate of CD4 decline was determined using a linear mixed effect model (LME). The rate of CD4 decline between the R5 and X4 groups was compared using a two-tailed Mann-Whitney test. *P* value < 0.05 is considered to be statistically significant.

### The “escape by shifting concept: Viral entry tropism shift as an evolutionary mechanism of immune evasion

Our study supports a novel concept that HIV-1 coreceptor switch is an evolutionary mechanism of immune evasion. This is based on multiple lines of evidence. First, the mutations responsible for coreceptor switch were strongly selected for at the peak of autologous neutralizing activity and conferred escape to neutralization. Second, both the N301 glycan site and V3 crown are common targets for V3 specific NAbs^27,34–36^. These NAbs have relatively high prevalence in HIV-1 infected people^36,41–45^, and their footprints overlap with the CCR5 binding region^27,33^. Third, the founder X4 viruses were immune escape variants with an impaired ability to use CCR5. Fourth, phenotypically confirmed X4 viruses from different HIV-1 subtypes, including those investigated here and in previous studies, are in general poorly neutralized by V3 NAbs^27,28,46^. These findings taken together support the following scenario underlying the phenomenon of coreceptor switch: During the co-evolution of HIV-1 and host immune responses, certain variants exploit genetic pathways to evade autologous neutralization at the cost of CCR5 tropism (due to the overlap between the CCR5 binding region and certain antigenic epitopes). In this case, mutation(s) which can confer CXCR4 usage will compensate for the impairment of CCR5-mediated entry, and thus be selected for in the viral quasispecies.

Indeed, the overlap between virus antigenic epitopes and receptor binding sites is widely observed for viruses from diverse families, and emerging evidence suggests that many viruses have the potential to use more than one receptor or coreceptor^47,48^. Potential co-evolution of virus antigenicity and receptor specificity has been observed for other viruses *in vitro*^49,50^. These lines of evidence prompted us to propose a general conceptual model: “escape by shifting” (Extended Data Fig. 10). The central hypothesis is that for viruses with receptor flexibility (for simplification, we use the term “receptor” to refer both receptor and coreceptor), entry tropism alteration functions as an evolutionary mechanism of immune evasion in the natural host with adaptive immunity, when the antigenic epitope overlaps with the receptor binding region. We coin the term “receptor tropism space” to describe the repertoire of receptors that could be utilized by a particular virus. Under immune pressure, a virus explores its “receptor tropism space” while exploring the sequence space and the fitness landscape. Immune escape mutations and compensatory mutations could both confer novel receptor usage, which functions as a mechanism to maintain virus entry capacity while evading host immune recognition. During this process, certain variants can achieve immune escape by altering the receptor they use. Consequently, the target cell specificity, pathogenicity, transmissibility, and the host range could change.

## Discussion

Identification of the co-evolution of HIV-1 antigenicity, coreceptor usage and CD4 subset targeting in natural HIV-1 infection advances our understanding of the immunopathogenesis of HIV-1. The poor neutralization susceptibility of X4 viruses to V3 specific bNAbs has previously raised the possibility that humoral response could select for X4 viruses^27,28,46^. However, major knowledge gaps have remained, including how the X4 variants originated *in vivo* and why they tend to be resistant to certain NAbs. By tracking the natural evolutionary pathway of the T/F virus, we identified the exact mutations responsible for both coreceptor switch and immune escape. Based on the “escape by shifting concept” we proposed, the switch to CXCR4 tropism represents a particular mechanism of fitness compensation at the entry level upon immune escape. In this regard, it would be necessary to understand the impact of immune escape mutations on HIV pathogenesis in a more comprehensive way, in particular their impact on the specificity of target cells, including the macrophages which could not be infected efficiently by the T/F HIV-1^51,52^.

The precise identification of the earliest moment of coreceptor switch allowed us to demonstrate that X4 phenotype is a causal factor of accelerated CD4 loss. It is generally considered that X4 viruses are more virulent because they can productively infect a broader range of CD4+ T cells, including the naïve subset which is important for CD4 homeostasis and renewal^9,53,54^. Our study provides novel insights into the pathogenesis of X4 tropic HIV-1. While the X4 viruses had a replication advantage in the naïve and the CM subsets, they could not outcompete the R5 viruses in the EM and TM subsets. The accelerated CD4 decline upon coreceptor switch was mainly due to the loss of the naïve and the CM cells. These observations together provide direct evidence that an altered, rather than expanded CD4 subset targeting could be a key factor in driving the loss of CD4 homeostasis. Several mechanisms could be involved. First, productive infection can kill the cells directly^55,56^. Second, considering the relatively quiescent or resting status of the naïve and CM cells, it is reasonable to hypothesize that a higher proportion of the cells could be abortively infected than those infected productively. Abortive infections can induce pyroptosis, which could in turn enhance immune activation and inflammation^57^. Third, the migration of the X4 viruses into the naïve and the CM cells would increase the virus burden in lymph nodes due to the expression of the homing receptor CCR7 on these cells^39,58^. While the exact mechanisms remain to be determined, these observations indicate the importance of CD4 subset preference in influencing the CD4 virulence of HIV-1. Indeed, preferential sparing of the CM CD4 subset is one of the potential mechanisms for the nonprogressive nature of simian immunodeficiency viruses (SIVs) infection in their natural hosts^59–61^. In HIV-1 infected people, a lower infection level of CM cells is a feature of long-term nonprogressors^62^. In light of this, it will be necessary to determine the potential heterogenicity of CD4 subset targeting of the R5 HIV-1. It is well known that R5 HIV-1 are heterogeneous in pathogenicity, and R5 viruses isolated from advanced disease could be more virulent than those isolated earlier from the same individual^19,63–65^. Whether this could be due to an enhanced ability to directly infect the CM cells requires further investigation.

The different CD4 subset preferences between R5 and X4 HIV-1 also sheds new light on understanding their distinct transmissibility. It has been a long-term question as to why R5 viruses are preferentially transmitted while a higher proportion of CD4+ T cells express CXCR4. Our study demonstrates a clear replication advantage of the R5 viruses in the EM and TM CD4 subsets when co-existing with the X4 viruses *in vivo*. Because the EM cells are more abundant in the mucosal tissues than the CM and the naïve CD4+ T cells^58,66,67^, a replication advantage in these cells is expected to provide an advantage during mucosal transmission. This is supported by an *ex vivo* study using the cervicovaginal explant model, which showed that the majority of CD4+ T cells in the tissues exhibited EM phenotype and preferentially supported the infection of R5 virus^68^. Moreover, in comparison to the naïve and CM cells, the EM and TM cells are likely to have a higher viral burst size due to their more activated status, and thus release more virions into the plasma. Indeed, the R5 viruses remained predominant in plasma in all participants harboring X4 variants in the current study. This could be another factor in determining the transmission advantage of R5 tropic HIV-1. In contrast, the altered tropism of the X4 HIV-1 towards the naïve and CM cells may compromise the transmissibility while enhancing virulence. These together also indicate that the CD4 subset preference could be a biological mechanism in governing the transmission-virulence tradeoff of HIV-1.

It has become increasingly apparent that many viruses have the potential to use more than one receptor or coreceptor, including the recently emerged SARS-CoV2^47,48,69^. The escape by shifting concept we proposed here is not specific for HIV, but a general concept for any virus with receptor flexibility. While potential co-evolution of virus antigenicity and receptor specificity has been observed for other viruses *in vitro*, such as the foot-and-mouth disease virus^49,50^, evidence from natural viral infections remains lacking. Like the CCR5 coreceptor for HIV, it is possible that a primary receptor is needed for a virus to establish efficient transmission in a particular host. The escape variants with altered receptor specificity could lose transmission advantage, or migrate into cells with smaller burst size, and consequently become less visible or difficult to identify by regular sequencing approaches. However, they could influence disease outcome. While the within-host quasispecies complexity is well documented for many RNA viruses, future research focusing on the within-host phenotype diversity, especially the potential entry tropism diversity, may provide novel insights into viral pathogenesis, as well as therapeutic and prevention strategies. It may also shed new light on how viruses gain zoonotic potential in the natural reservoir species.

## Figure Legends

**Extended Data Fig. 1.**
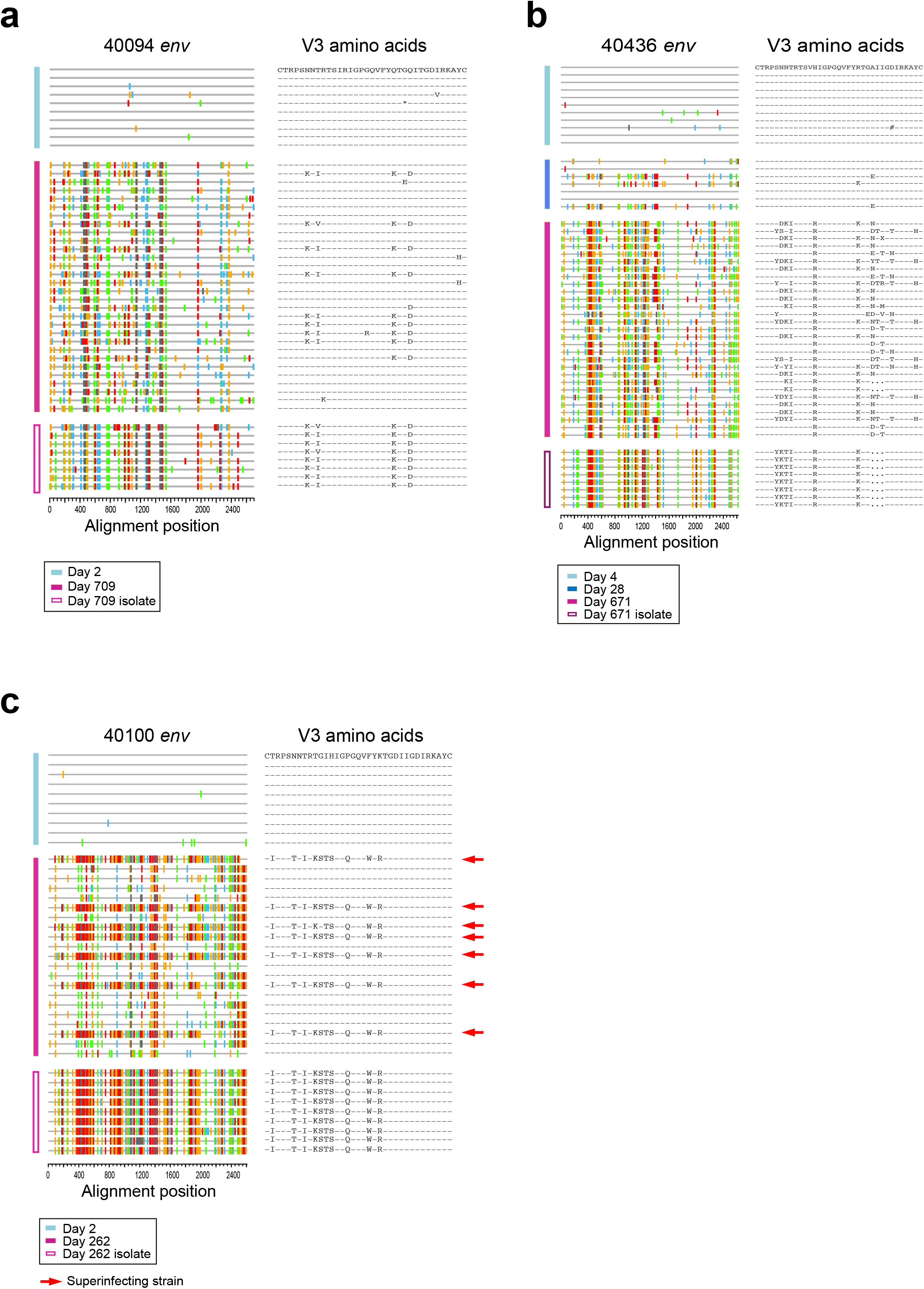
Genetic relationship between the X4-using viral isolates and the plasma viruses. **a-c**, The SGA sequences of the X4-using isolates are aligned to the SGA sequences of the plasma viruses. Sequences from different time points are color coded. Participant 40100 was initially infected by a single T/F virus and was super-infected by an X4 strain at day 262. The sequences of the superinfecting strain are indicated by red arrows. The sequences of the X4-using isolate matched the super-infecting strain.

**Extended Data Fig. 2.**
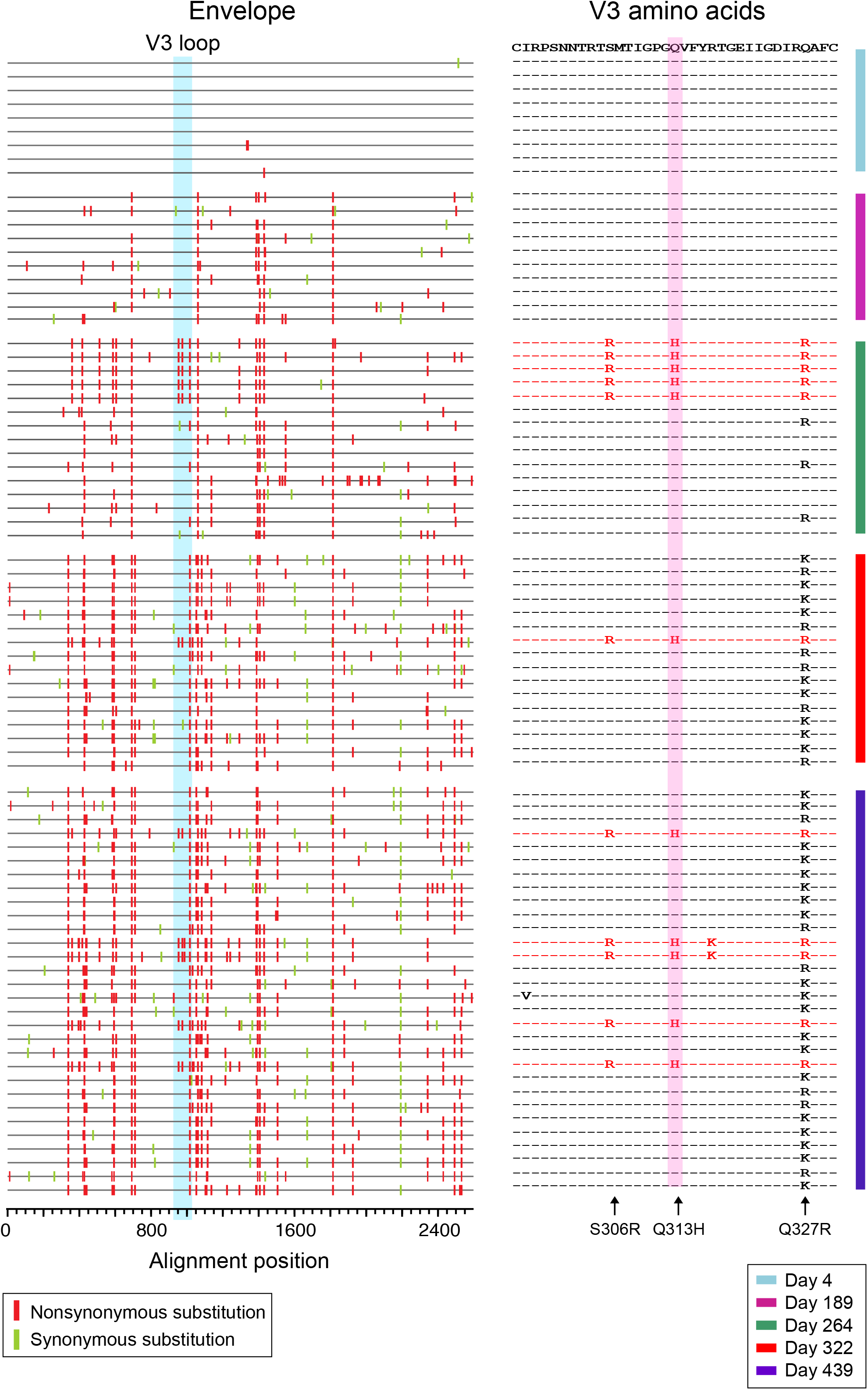
The evolutionary pathway leading to coreceptor switch of the 40257 T/F virus. Longitudinal envelope sequences of participant 40257 were obtained by SGA. The longitudinal evolution of the T/F virus is illustrated by using the highlighter plot. Sequences from different time points are color coded. The sequence on the top represents the T/F virus. The red and green tics indicate non-synonymous and synonymous substitutions compared to the T/F virus, respectively. The V3 loop is highlighted in blue. In the V3 amino acid alignment, the mutation responsible for coreceptor switch is shaded in red.

**Extended Data Fig. 3.**
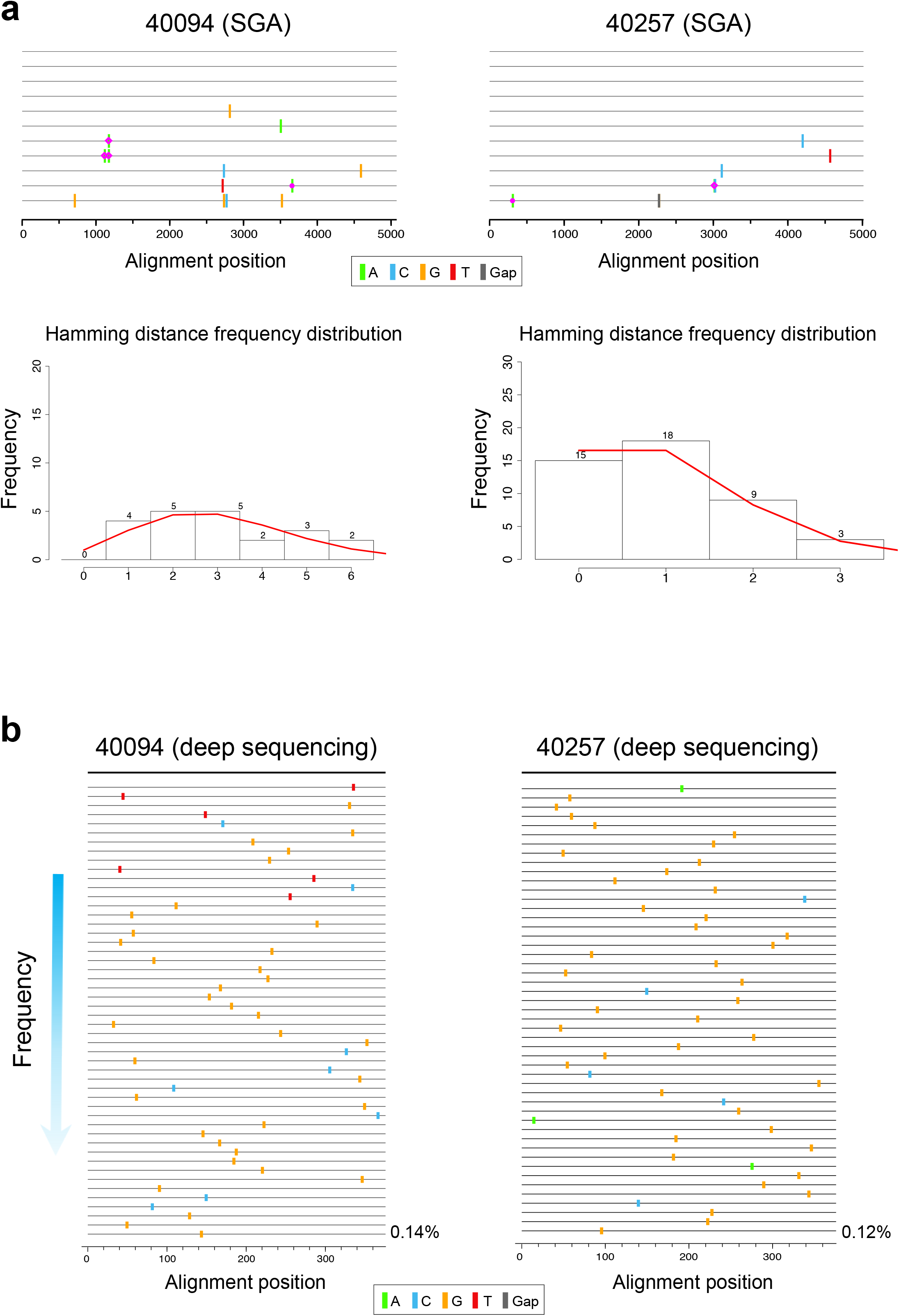
Determination of single T/F virus transmission in participants 40094 and 40257. **a**, SGA sequences obtained from acute infection were used to determine the number of T/F virus by using the Poisson-Fitter tool in the Los Alamos HIV Sequence Database. For both participants, the Hamming Distance (HD) frequency distributions follow Poisson distribution, and the sequences follow a star-phylogeny. **b**, Deep sequencing of the C2V3 region detected a single viral lineage during acute infection. The sequences are ordered according to their frequency. A total of 50 unique sequences are shown. The T/F sequence is used as the master sequence (the black line on the top). The frequency of the last sequence in the highlighter plot is shown.

**Extended Data Fig. 4.**
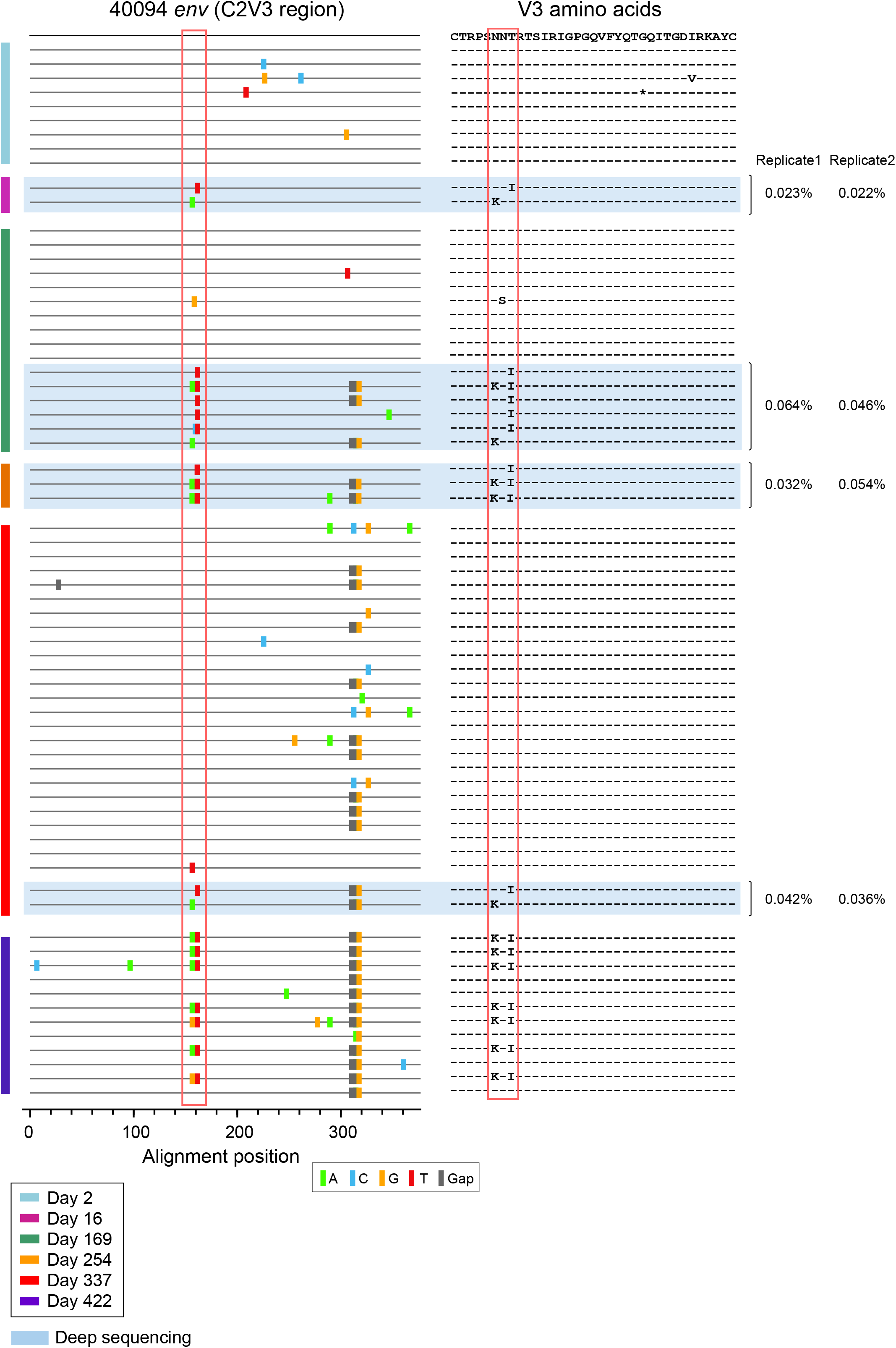
Evolution of the viral variants carrying the driver mutations in participant 40094. The deep sequencing reads carrying the driver mutations (shaded in blue) are aligned to plasma viral sequences obtained by SGA. Sequences from different time points are color coded. The mutations responsible for coreceptor switch are indicated in red boxes. Deep sequencing was carried out in duplicate. The frequency of the deep sequencing reads is shown on the right.

**Extended Data Fig. 5.**
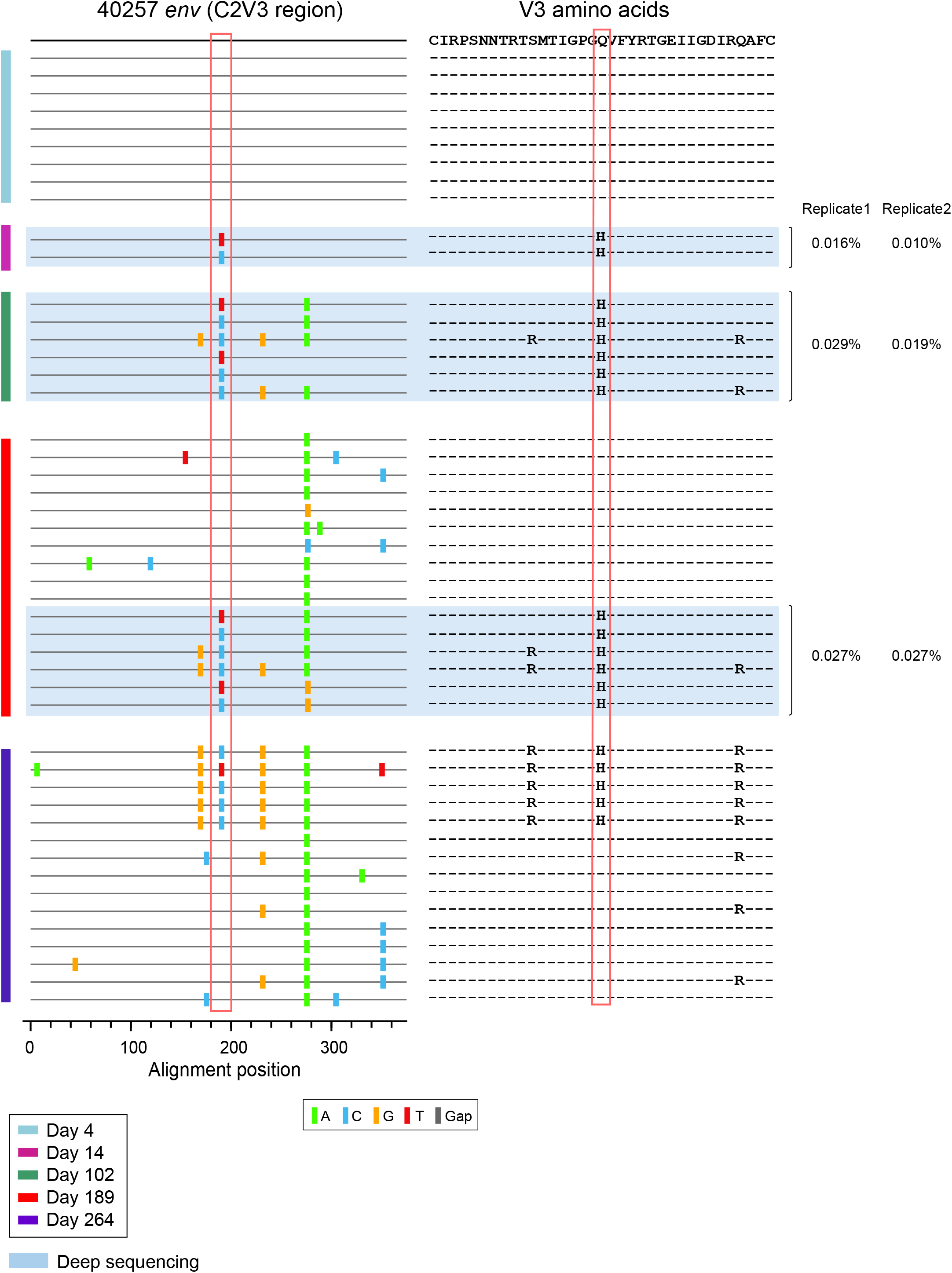
Evolution of the viral variants carrying the driver mutations in participant 40257. The deep sequencing reads carrying the driver mutations (shaded in blue) are aligned with plasma viral sequences obtained SGA. Sequences from different time points are color coded. The mutations responsible for coreceptor switch are indicated in red boxes. Deep sequencing was carried out in duplicate. The frequency of the deep sequencing reads is shown on the right.

**Extended Data Fig. 6.**
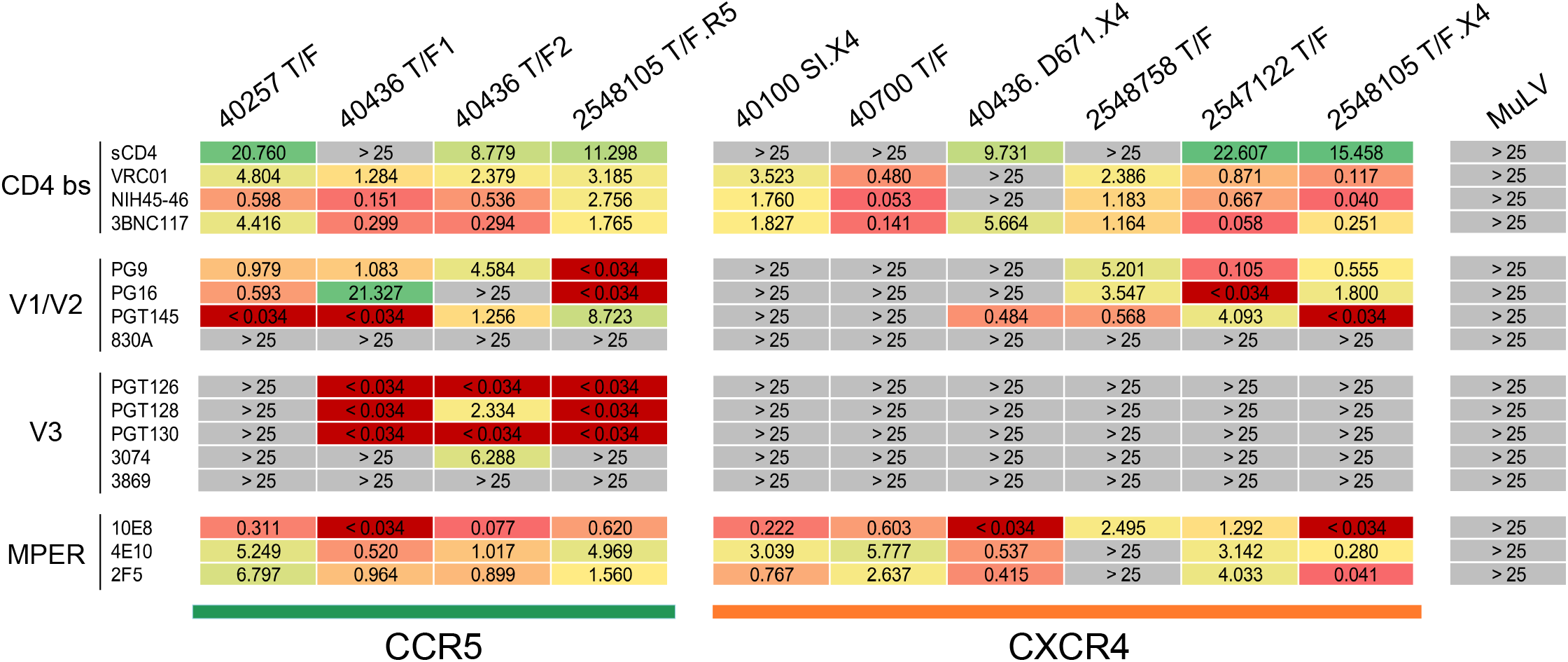
Neutralization sensitivity of R5 and X4 HIV-1 to broadly neutralizing antibodies (bNAbs). The CCR5 group includes four R5 tropic T/F viruses identified in the RV217 and RV254 cohorts. The CXCR4 group includes six phenotypically confirmed X4-using viruses, including the superinfecting strain in 40100, the X4-tropic T/F virus in 40700, an X4 variant isolated from 40436, as well as three X4-using T/F viruses identified in participants 2548758, 2547122 and 2548105. Among them, the 40700 T/F virus uses CXCR4 exclusively, while the other five viruses have low-level R5 using ability. The IC_50_ (µg/mL) of each bNAb is shown. The MuLV was used as the negative control.

**Extended Data Fig. 7.**
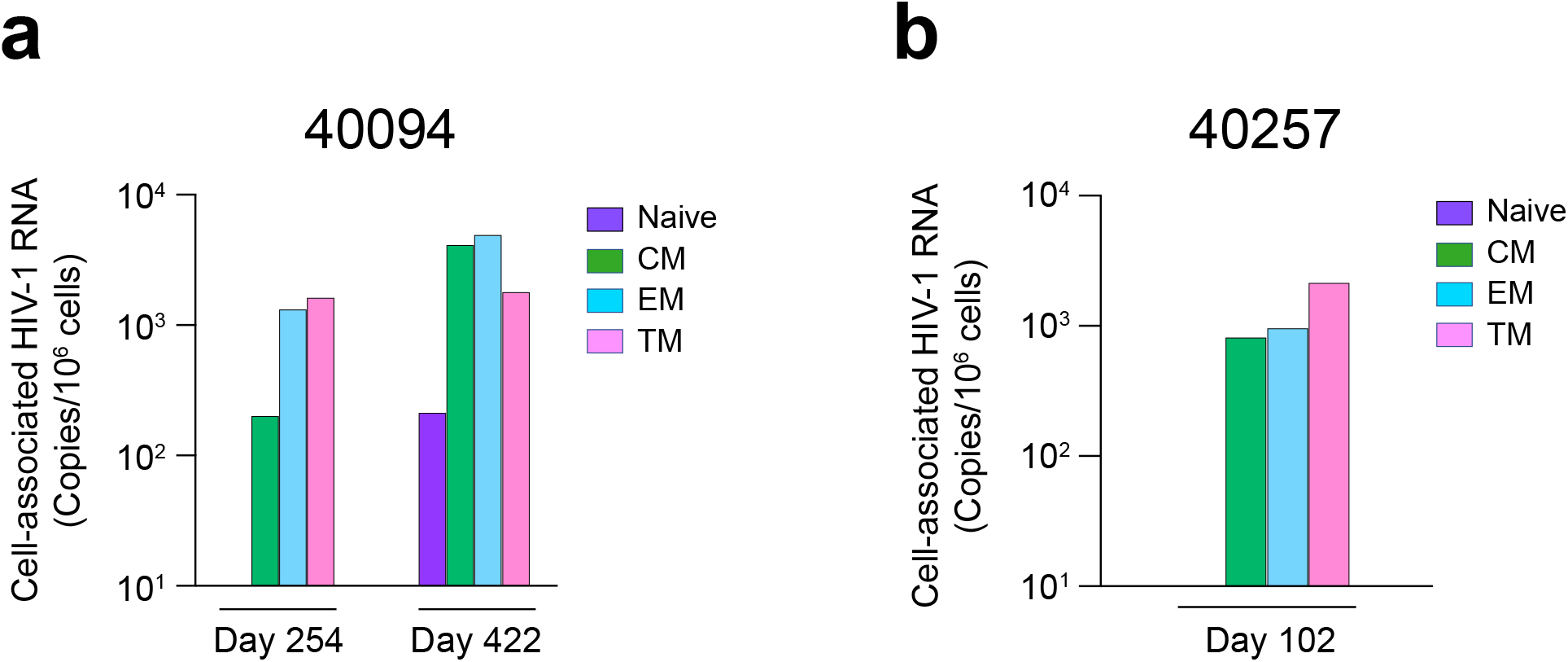
Quantification of cell associated HIV-1 RNA in participants 40094 and 40257. **a**, Cell associated HIV-1 RNA in different CD4 subsets were quantified in participant 40094 at day 254 (before coreceptor switch) and day 422 (the earliest time point of coreceptor switch). The cell associated RNA was undatable in the naïve subset at day 254. **b**, Cell associated HIV-1 RNA in different CD4 subset were determined in participant 40257 at day 102 (before coreceptor switch). The cell associated RNA was undetectable in the naïve subset.

**Extended Data Fig. 8.**
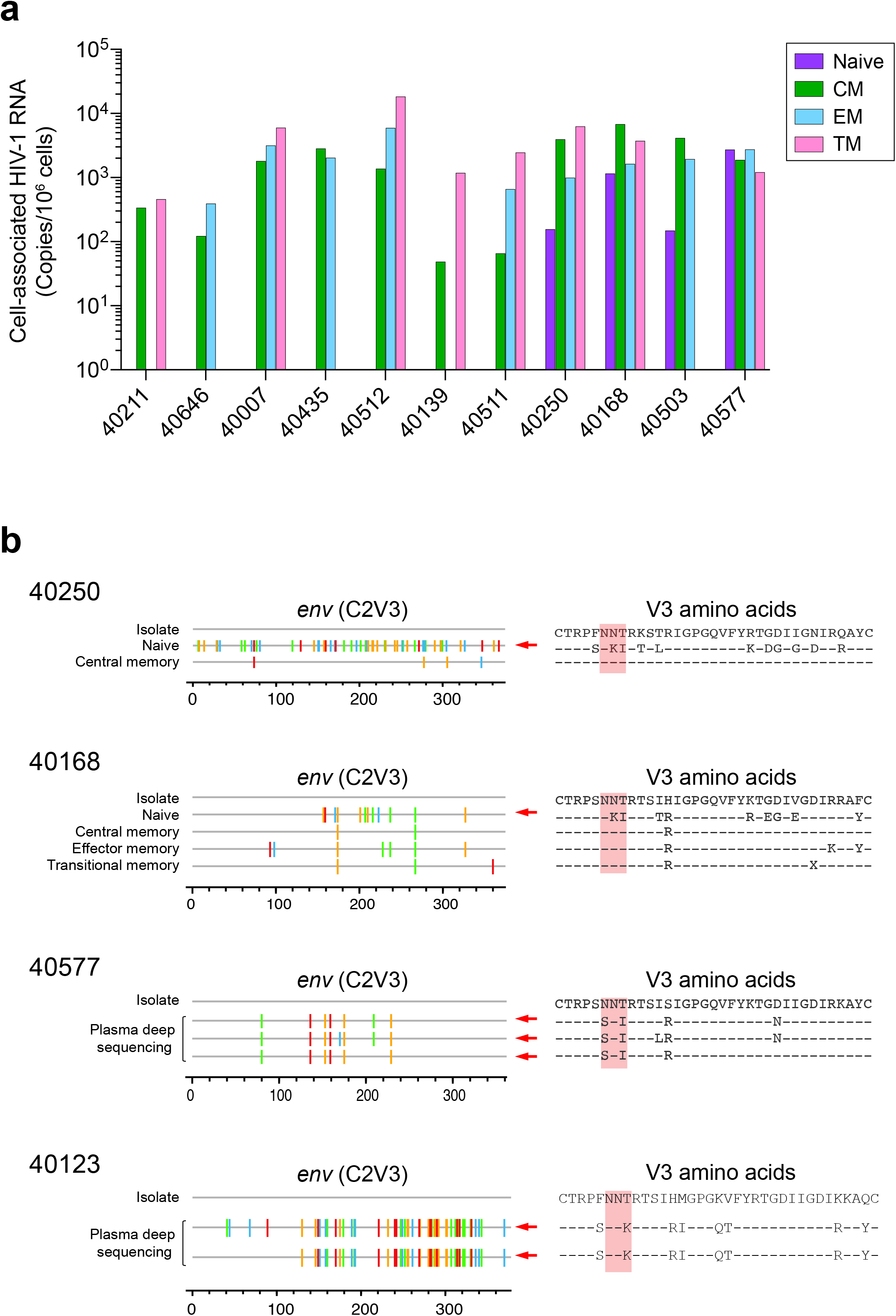
Detection of X4 variants in naïve CD4+ T cells or in plasma in participants from whom only R5 viruses were isolated. **a**, Cell associated HIV-1 RNA in different CD4 subsets of 11 participants for whom the primary viral isolates showed pure R5 phenotype. **b**, X4 viruses were detected in naïve CD4+ T cells for participants 40250 and 40168. A low frequency of X4 variants was detected in plasma in participants 40577 and 40123 by deep sequencing. The phenotypically confirmed X4 variants are indicated by red arrows.

**Extended Data Fig. 9.**
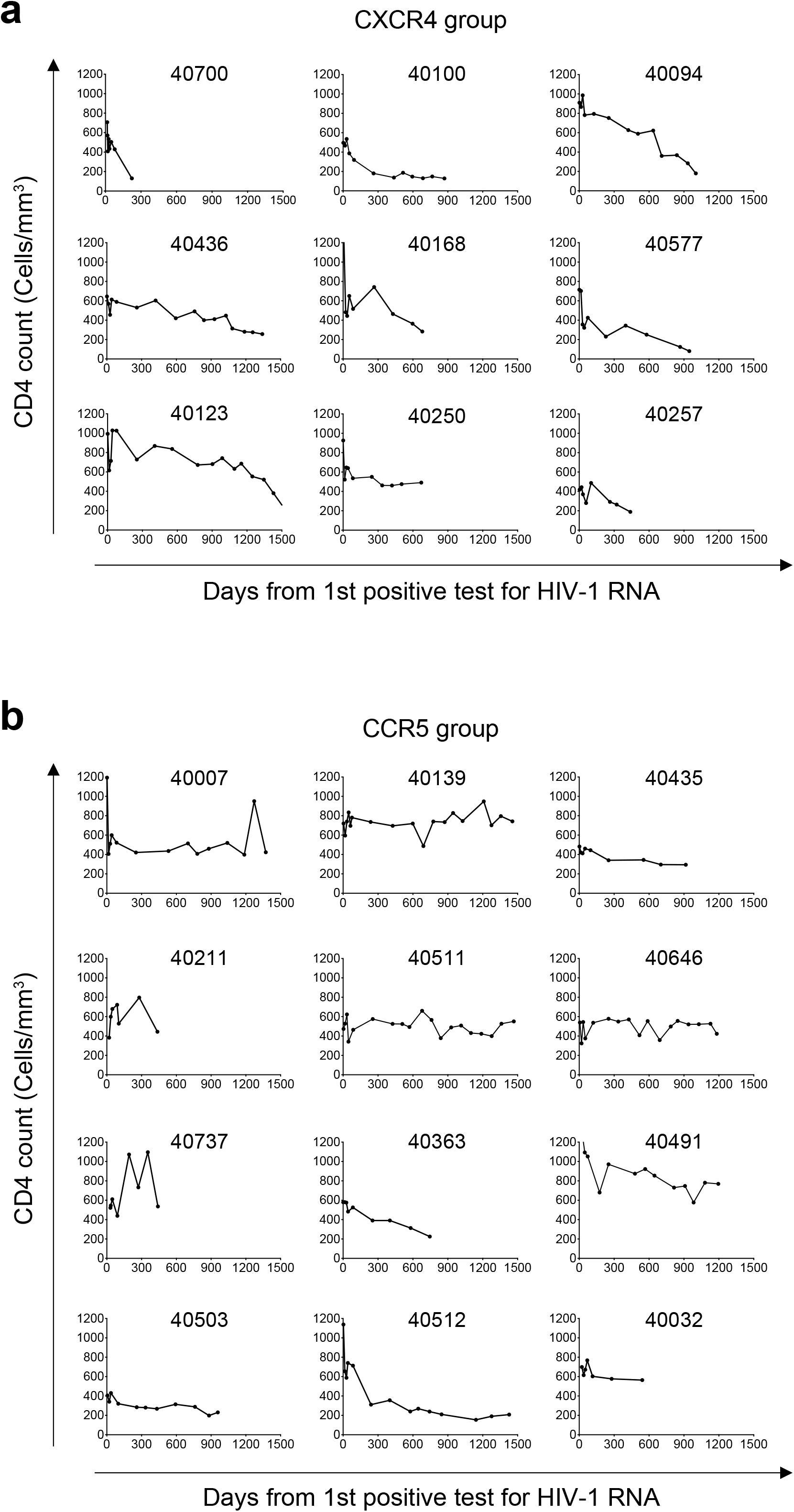
CD4 dynamics of each study participant. **a**, CD4 dynamics of participants harboring X4 viruses. Participant 40700 was infected by an X4 T/F HIV-1 without CCR5 using ability. **b**, CD4 dynamics of participants without the evidence of harboring X4 variants. The longitudinal data from the earliest available time point to the last available time point before ART initiation is shown.

**Extended Data Fig. 10.**
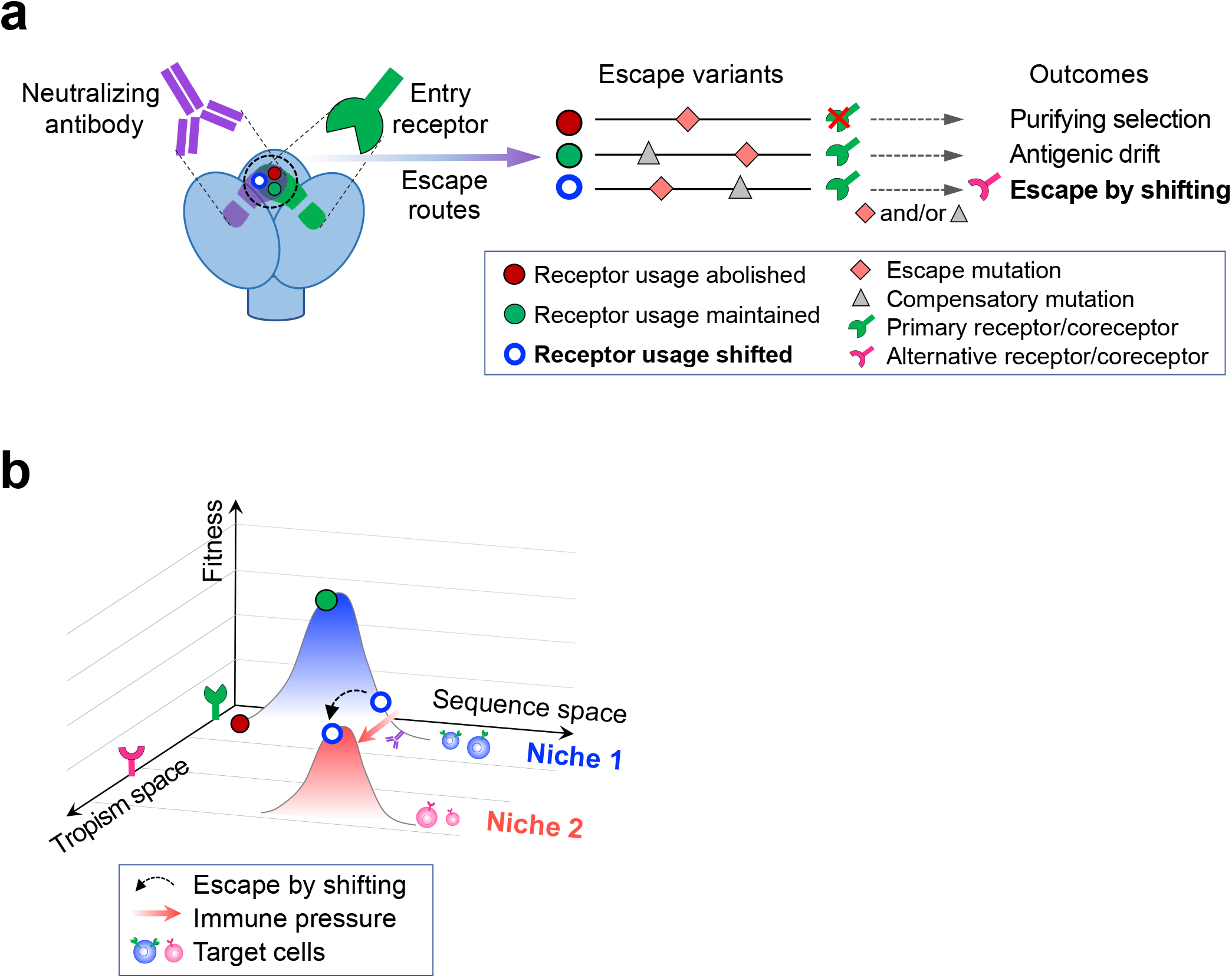
Conceptual framework of the “escape by shifting” concept. **a**, When the footprint of a neutralizing antibody overlaps with virus receptor binding site, receptor usage alteration functions as an evolutionary route of immune evasion. **b**, Escape variants explore receptor tropism space, sequence space and fitness landscape under host immune pressure. Alteration in receptor usage allows an escape variant to find its fitness peak in a novel niche.

## Methods

### Study participants

Study participants were from the RV217 and the RV254 cohorts who were identified very early in acute HIV-1 infection^32,70^ (Supplementary Table 1). PBMC and plasma samples of 21 participants in the RV217 Thailand cohort^32^ who had pre-ART samples available up to approximately two years from HIV-1 transmission were used in the current study. Four phenotypically confirmed T/F *env* clones, including one R5 tropic and three X4 tropic, were identified in participants of the RV254 cohort who had acute HIV-1 sequences generated for previous studies^71,72^. All study participants were infected by CRF01_AE HIV-1 except for participants 40168 and 2548758 who were infected by CRF01_AE/B recombinant viruses (Supplementary Table 1). Written consent was provided by all participants. The study was approved by the local ethics review boards, the Walter Reed Army Institute of Research, and the institutional review boards of the University of Maryland School of Medicine.

### Virus isolation

To isolate viruses from HIV-1 infected individuals, PBMCs from a healthy donor were stimulated for 3 days in RPMI1640 containing 10% fetal bovine serum (FBS), interleukin 2 (IL-2) (50 U/ml; PeproTech), soluble anti-CD3 (0.5 μg/ml; eBioscience) and soluble anti-CD28 (0.5 μg/ml; eBioscience). After stimulation, the cells were washed twice with RPMI1640 to remove the antibodies. A total of 10^7^ stimulated PBMCs from a healthy donor were then mixed with 10^7^ of cryopreserved PBMCs from an HIV-1 infected individual. The cells were cultured in a T25 flask in RPMI1640 containing 10% FBS and 50 U/ml IL-2 for up to 4 weeks. Every 3 days, half volume of the culture supernatant was replaced with fresh medium. Every 7 days, half of the culture (including the cells) was removed, and a total of 5 × 10^6^ of stimulated PBMCs from a healthy donor were added. The p24 concentration in the culture supernatant was measured every week. The viruses were harvested between two and four weeks when the p24 concentration in the culture supernatant achieved at least 5 ng/ml.

### Single genome amplification

Single genome amplification (SGA) was performed as previously described^73^. In brief, HIV-1 RNA in plasma or in the primary viral isolates was extracted using the QIAamp Viral RNA Mini Kit (Qiagen). cDNA was synthesized using the SuperScript III reverse transcriptase (Invitrogen) using the primer 1.R3.B3R 5’-ACTACTTGAAGCACTCAAGGCAAGCTTTATTG-3’ (nt 9642-9611 in HXB2). To amplify the 3’ half viral genome, the first round PCR was performed using the primers 07For7 5’-CAAATTAYAAAAATTCAAAATTTTCGGGTTTATTACAG-3’ (nt 4875-4912) and 2.R3.B6R 5’-TGAAGCACTCAAGGCAAGCTTTATTGAGGC-3’ (nt 9636–9607), and the second round PCR was performed with the primers VIF1 5’-GGGTTTATTACAGGGACAGCAGAG-3’ (nt 4900-4923) and Low2c 5’-TGAGGCTTAAGCAGTGGGTTCC-3’ (nt 9591-9612). Two microliters of the first round PCR products were used for the second round PCR amplification. The PCR thermocycling conditions were as follows: one cycle at 94°C for 2 min; 35 cycles of a denaturing step at 94°C for 15 sec, an annealing step at 60°C for 30 sec, an extension step at 68°C for 4 min; and one cycle of an additional extension at 68°C for 10 min. The PCR amplicons were directly sequenced by the cycle sequencing and dye terminator methods. Individual sequences were assembled and edited using Sequencher (Gene Codes). The sequences were aligned using the Gene Cutter in the Los Alamos HIV Sequence Database, followed by manual adjustment to obtain optimal alignment.

### Pseudovirus preparation and titration

To generate HIV-1 envelope (*env*) clones, the *rev-vpu-env* cassette was PCR amplified from the corresponding SGA products or chemically synthesized (GenScrip) and cloned into the expressing vector pcDNA3.3-TOPO (Invitrogen). The desired mutations were introduced by site-directed mutagenesis. All envelope clones were confirmed by sequencing. The pseudovirus stocks were prepared as previously described^74^. In brief, 2 μg of *env* clone was co-transfected with 4 μg of the pNL4.3-ΔEnv-vpr+-luc+ into 293T cells in a T25 flask using the FuGENE6 transfection reagent (Promega). The cells were cultured at 37°C for 6 hours before the medium was completely replaced with fresh medium. The culture supernatants containing the pseudoviruses were harvested at 72 hours post transfection, aliquoted and stored at −80°C until use. The infectious titers (TCID_50_) of the pseudovirus stocks were determined on TZM-bl cells.

### Determination of coreceptor usage

Coreceptor usage of the primary viral isolates and the pseudovirus were determined on NP-2 cells lines expressing CD4 together with either the CCR5 or CXCR4 coreceptors as previously described^74^. The parental NP-2 cell line expressing CD4 alone was used as a control. To determine the coreceptor usage of primary viral isolates, NP-2 cell lines were seeded in a 48-well plate at a density of 2 × 10^5^ cells per well one day before infection. The next day, the cells were infected with 200 µL undiluted viral isolates. After 6 hours incubation at 37°C, the cells were washed 3 times and cultured in fresh medium at 37°C for 72 hours. The p24 concentration in the culture supernatant was measured three days post infection (PerkinElmer). Virus infectivity was considered positive if the p24 concentration was at least three times higher than the background concentration in parental cell line. All infections were performed in duplicates.

To determine the coreceptor suage of the pseudovirus, NP-2 cells were seeded in a 96-well plate one day before infection at a density of 1 × 10^5^ cells per well. On the next day, the cells were infected with approximately 200 TCID_50_ of each pseudovirus (MOI = 0.002). After 6 hours of incubation at 37°C, the infected cells were washed twice with the culture medium and cultured at 37°C for three days. At 72 hours post infection, the infected cells were lysed, and the infectivity was determined by measuring the relative luciferase units (RLU) in the cell lysates using the Britelite plus system (PerkinElmer). Viral infectivity was considered positive if the RLU value was at least 5-fold higher than the background RLU value in the parental cell line. All experiments were performed in triplicates.

### Next generation sequencing

The library for sequencing on Illumina MiSeq was prepared using a nested PCR approach as described previously^25^. The first round PCR was carried out using the forward primer 44F 5’-ACAGTRCARTGYACACATGG-3’ (nt 6954-6973) and the reverse primer 35R 5’-CACTTCTCCAATTGTCCITCA-3’ (nt 7648-7668). The first round PCR conditions were as follow: one cycle at 94°C for 2 min; 30 cycles of a denaturing step at 94°C for 15 sec, an annealing step at 55°C for 30 sec, an extension step at 68°C for 1 min; and one cycle of an additional extension at 68°C for 10 min. A total of 5 µL of first round PCR products were used for second round PCR amplification. The second round PCR was carried out using a panel of primers containing the Illumina index and adaptors as previously described^25^, using the following PCR conditions: one cycle at 94°C for 2 min; 5 cycles of a denaturing step at 94°C for 15 sec, an annealing step at 55°C for 30 sec, an extension step at 68°C for 1 min; and another 30 cycles of a denaturing step at 94°C for 15 sec, an annealing step at 60°C for 30 sec, an extension step at 68°C for 1 min; and one cycle of an additional extension at 68°C for 10 min. The second round PCR products were gel purified, and the final library was generated by pooling the second round PCR product from each sample with an equal concentration. To increase the diversity of the library, 40% of the PhiX was spiked in (Illumina). The library was sequenced on the Illumina MiSeq using the MiSeq Reagent Kit v3 (Illumina). Deep sequencing was performed in duplicate for each sample.

### Next generation sequencing data analysis

The raw fastq reads in files “read 1” and “read 2” were merged using the FLASH software^75^. The merged fastq files were then filtered based on data quality using the following parameters: no more than 10 base calls with Q score lower than 30 in each read. After filtering, identical reads in each sample were collapsed to generate the unique sequences. Singletons (sequences which only appeared once in a sample) were excluded from the downstream analysis. To minimize the possibility that a unique sequence was generated due to sequencing error, deep sequencing was performed in duplicate for each sample. Only the deep sequencing reads that appeared in both experiments were used for analysis. The deep sequencing data was processed and analyzed using publicly available software and tools on the Galaxy platform^76^.

### Neutralization assay

The neutralization activity of plasma samples and monoclonal antibodies (mAbs) was measured by using a luciferase reporter system in TZM-bl cells. Plasma samples were heat inactivated at 56°C for 45 minutes. The inactivated plasma was diluted at a 1:3 serial dilution starting from 1:20. The mAbs were diluted at a 1:3 serial dilution from a starting concentration of 25 μg/mL. The virus stocks were diluted to a concentration that achieved approximately 150,000 RLU in the TZM-bl cells (or at least 10 times above the background RLU of the cells control). The serial diluted plasma samples or mAbs were then incubated with the viruses for 1 hour at 37°C in duplicate before the TZM-bl cells were added. The 50% inhibitory dose (ID_50_) was determined as the dilution at which the relative luminescence units (RLUs) were reduced by 50% in comparison to the RLUs in the virus control wells after subtraction of the background RLUs in cell control wells.

### CD4 subset analysis and sorting

PBMCs from HIV-1 infected individuals were stained by the following antibodies: CD3-Brilliant Violet 605 (clone OKT3, BioLegend), CD4-PerCP-Cy5.5 (clone OKT4, Biolegend), CCR7-PE-CF594 (clone 2-L1-A, BD Biosciences), CD27-PE (clone M-T271, BioLegend), CD45RO-APC (clone UCHL1, BioLegend). The cells were then stained by the Live-dead aqua prior to flow analysis to exclude the dead cells (Invitrogen). The stained cells were sorted on a BD FACSAria II cell sorter (BD Biosciences). Four CD4 subsets were defined as follows: naïve (CD45RO^-^, CCR7^+^, and CD27^+^), central memory (CD45RO^+^, CCR7^+^, and CD27^+^), transitional memory (CD45RO^+^, CCR7^-^, and CD27^+^) and effector memory (CD45RO^+^, CCR7^-^, and CD27^-^). The purity of each sorted subset was higher than 95%.

### Determination of CCR5 and CXCR4 expression

To determine the level of CCR5 and CXCR4 expression on each CD4 subset, PBMCs were stained with the following antibodies: CD3-Brilliant Violet 605 (clone OKT3, BioLegend), CD4-PerCP-Cy5.5 (clone OKT4, BioLegend), CCR7-PE-CF594 (clone 2-L1-A, BD Biosciences), CD27-FITC (clone M-T271, BioLegend), CD45RO-APC (clone UCHL1, BioLegend), CCR5-PE (clone J418F1, BioLegend) or CXCR4-PE (clone Q18A64, BioLegend). The CCR5 and CXCR4 staining antibodies were titrated to determine the optimal concentration. To evaluate the non-specific staining, fluorescent stanning was cold-inhibited by a 100-fold excess of the unlabeled CCR5 or CXCR4 antibody (the same clone as the labeled antibody) mixed with the respective labeled antibody. A fluorescence minus one (FMO) staining was also determined for CCR5/CXCR4 staining. The highest concentration of the labeled antibody with which the cold inhibition showed virtually overlapping staining with the FMO was used to quantify the levels of CCR5 and CXCR4 expression on each CD4 subset. All flow data were collected using the DIVA 7.0 software on the FACSAria II (BD Biosciences) cell sorter/analyzer and analyzed by the FlowJo software (FlowJo LLC, Ashland, OR).

### Quantification of cell associated HIV-1 RNA

Cell associated HIV-1 RNA was quantified for each sorted CD4 subset by amplifying part of the *pol* gene. RNA was extracted from the sorted cells using the RNeasy Mini kit (Qiagen). A total of 8.5 µL extracted RNA was subjected to one-step RT-PCR using the Superscript III one-step RT-PCR system (Invitrogen). The one-step RT-PCR was performed using the forward primer Pol F1 5’-TACAGTGCAGGGGAAAGAATA-3’ (nt 4809-4829) and the reverse primer Pol R1 5’-CTTCTTGGCACTACTTTTATGTCAC-3’ (nt 4993-5017). The PCR conditions were as follow: a reverse transcription step at 50°C for 1h; A denaturing step at 94°C for 2 min; 16 cycles of a denaturing step at 94°C for 15 sec, an annealing step at 55°C for 30 sec, an extension step at 68°C for 1 min, and one cycle of an additional extension at 68°C for 5 min. The first round PCR products were diluted 10-fold and a total of 6.4 µL of diluted PCR products were used for the real-time PCR using the forward primer Pol F1, the reverse primer Pol R2 5’-CTGCCCCTTCACCTTTCC-3’ (nt 4957-4974), and the probe Pol Famzen: 5’-/56-FAM/TTTCGGGTT/ZEN/TATTACAGGGACAGCAG/3IABkFQ/-3’ (nt 4896-4921). The real-time PCR was performed on the QuantStudio 3 Real-Time PCR Systems (Thermo Fisher Scientific) using the following conditions: A denaturing step at 94°C for 4 min, 45 cycles of a denaturing step at 94°C for 3 sec, an annealing and extension step at 60°C for 20 sec. The copy number of the input RNA was determined by using the RNA standard generated by *in vitro* transcription. In brief, the amplicon region (we used the CRF01_AE consensus sequence in this study) was cloned into the pUC57 vector downstream of the T7 promoter. The DNA fragment containing the amplicon was PCR amplified and the RNA was generated by *in vitro* transcription using the MEGAscript T7 Transcription Kit (Invitrogen).

### Determination of the rate of CD4 decline

The rate of CD4 decline was determined using a linear mixed effect model (LME)^77^. The LME model was hierarchical in the sense that it estimated a population specific slope and intercept with time, as well as subject-specific slopes and intercepts. The longitudinal data contained CD4 data from the earliest available time point to the last available time point before ART initiation was used for the analysis. The rate of CD4 decline between the R5 and X4 groups was compared using a two-tailed Mann-Whitney test.

## Acknowledgements

The authors thank the study participants of the RV217 and RV254/SEARCH 010 cohorts. We thank Sebastian Molnar and Amy Nguyen for technical assistance. This study was supported by the Institute of Human Virology, University of Maryland School of Medicine. Part of the study was supported by cooperative agreements between the Henry M. Jackson Foundation for the Advancement of Military Medicine, Inc., and the U.S. Department of Defense (DOD). RV254/SEARCH010 is supported by cooperative agreements (WW81XWH-18-2-0040) between the Henry M. Jackson Foundation for the Advancement of Military Medicine, Inc., and the United States Army Medical Research and Development Command (USAMRDC) and by an intramural grant from the Thai Red Cross AIDS Research Centre and, in part, by the Division of AIDS, the National Institute of Allergy and Infectious Diseases, National Institutes of Health (AAI21058-001-01000). Antiretroviral therapy for RV254/SEARCH 010 participants was supported by the Thai Government Pharmaceutical Organization, Gilead Sciences, Merck and ViiV Healthcare. M.H.M. and H.S. were supported by the NIH grant R21AI147893.

## Disclaimer

The views expressed are those of the authors and should not be construed to represent the positions of the U.S. Army, the Department of Defense, the National Institutes of Health, the Department of Health and Human Services, or the Henry M. Jackson Foundation for the Advancement of Military Medicine, Inc. The investigators have adhered to the policies for protection of human subjects as prescribed in AR-70-25.

## Author contributions

H.S., V.R.P., and M.H.M. designed the study. M.H.M., M.Z., L.W., E.S-B., M.B., A.M.O., F.D-M., S.S. and H.S. performed the experiments. M.H.M., E.S-B., M.B., A.M.O., S.T., and H.S. were involved in viral sequencing and genetic analysis. D.K. and L.F. modeled the CD4 dynamics. Y.T. contributed to the design and data analysis of the flow cytometry and cell sorting experiments. N.C. provided the protocol of cell associated HIV-1 RNA assay and contributed to discussions of the experiments and data. N.P., J.A., D.S., S.V., N.L.M., L.A.E., and M.L.R. were involved in recruiting of the clinical cohorts, analysis of the clinical data and supplied clinical samples. H.S. wrote the manuscript with the input from all authors. All authors read the manuscript and approved the submission.

## Data availability

The GenBank accession numbers of the sequences used in the current study, including newly generated sequences and sequences submitted previously were summarized in Supplementary Table 3.

